# Injection molded open microfluidic well plate inserts for user-friendly coculture and microscopy

**DOI:** 10.1101/709626

**Authors:** John H. Day, Tristan M. Nicholson, Xiaojing Su, Tammi L. van Neel, Ivor Clinton, Anbarasi Kothandapani, Jinwoo Lee, Max H. Greenberg, John K. Amory, Thomas J. Walsh, Charles H. Muller, Omar E. Franco, Colin R. Jefcoate, Susan E. Crawford, Joan S. Jorgensen, Ashleigh B. Theberge

## Abstract

Open microfluidic cell culture systems are powerful tools for interrogating biological mechanisms. We have previously presented a microscale cell culture system, based on spontaneous capillary flow of biocompatible hydrogels, that is integrated into a standard cell culture well plate, with flexible cell compartment geometries and easy pipet access. Here, we present two new injection molded open microfluidic devices that also easily insert into standard cell culture well plates and standard culture workflows, allowing seamless adoption by biomedical researchers. These platforms allow culture and study of soluble factor communication among multiple cell types, and the microscale dimensions are well-suited for rare primary cells. Unique advances include optimized evaporation control within the well, manufacture with reproducible and cost-effective rapid injection molding, and compatibility with sample preparation workflows for high resolution microscopy (following well-established coverslip mounting procedures). In this work, we present several use cases that highlight the usability and widespread utility of our platform including culture of limited primary testis cells from surgical patients, microscopy readouts including immunocytochemistry and single molecule fluorescence *in situ* hybridization (smFISH), and coculture to study interactions between adipocytes and prostate cancer cells.

## Introduction

An important goal of microscale cell culture systems is their translation and widespread adoption into everyday biomedical research.^1^ While the promise of microscale cell culture systems in biomedical research has been recognized for the past two decades, these technologies have only recently become well-poised for widespread adoption by biomedical researchers.^2,3^ ‘Open’ microfluidic devices, which contain channels with at least one air-liquid interface, have contributed to increased accessibility.^4^

Open microfluidics allows precise patterning of liquids and cell suspensions via spontaneous capillary flow.^5,6,7,8^ We have recently presented a 3D-printed well plate insert for cell culture, the Monorail Device, that utilizes spontaneous capillary flow to pattern biocompatible hydrogels on a surface, creating hydrogel walls that partition the well into separate chambers for cell culture.^9^ This platform enables a range of cell culture compartment geometries with physical partitioning of different cell types and the ability to study soluble factors exchanged in coculture through the hydrogel wall.^9^ Key advantages of this platform include compatibility with traditional cell culture platforms (e.g., well plates) so that cells can be grown on commercially available cell culture treated surfaces, ease of pipetting due to open microfluidic design, and the ability to pattern various shapes. Lee et al. presented a different platform based on similar principles, using injection molded polystyrene to create a 3D coculture system in the form of a 96-well plate; in this case, the entire well plate structure containing the fluidic features was manufactured as a single plastic structure, and the well plate floor was subsequently created by bonding adhesive tape to the injection molded structure.^10^

Both of these examples represent important advances in translating microscale cell-culture systems into formats that are easily utilized for biological applications. However, microscale cell culture platforms based on open and suspended microfluidics continue to have several challenges for cell culture applications that may limit widespread adoption by biolmedical researchers. These challenges include evaporation control at the air-liquid interface, variability from device fabrication and user operation, and difficulty interfacing with standard workflows for high resolution microscopy which involve culturing cells on coverslips and subsequent mounting on glass slides. Here, we present two new open microfluidic devices based on our previously established platform.^9^ These devices retain the advantages of the original iteration—easy integration with well plates that are familiar to biomedical researchers, flexible geometric patterning of biocompatible hydrogels, and pipet accessibility. Distinct advantages demonstrated in the present manuscript include simple and effective evaporation control strategies, manufacture with rapid injection molding, and compatibility with high resolution microscopy; these three considerations are reviewed in the following paragraphs.

Compared to conventional cell culture vessels such as flasks, petri dishes and well plates, microscale systems have a higher surface area to volume ratio, leading to less cell culture media per cell.^11^ The resulting cell stress can be mitigated by frequent media changes and decreased cell seeding density, but evaporation remains an important concern, and is of particular importance for microscale cell culture systems that are suited for rare, sensitive cell types affected by changes in osmolarity.^11,12,13,14^ A number of strategies have been employed to attenuate evaporation, including reservoirs of water on-chip, submersion of the entire device or container in water, and use of oil to cover aqueous liquids.^8,9,12^ In the present work, we present two approaches to evaporation control contained within the well plate that are conducive to cell viability and do not require placement of the entire well plate in a larger secondary evaporation control vessel.

Widespread adoption of microscale cell culture systems in biomedical applications is challenging because of the need for low cost production, reproducible manufacturing, and the ability to iterate on designs. Common methods for microfluidic device fabrication, which include micro-machining,^15^ soft lithography,^16^ hot embossing,^17^ and 3D printing,^18^ are better suited for early stage prototyping than mass production. Injection molding is the gold standard for mass manufacturing and offers high reproducibility and fast manufacturing times. Until recently, the downside of injection molding had been the high cost (up to tens of thousands of dollars) associated with producing complex high-quality steel molds. Rapid injection molding has recently lowered the initial mold cost significantly, and with this, microscale cell culture systems are now poised for high volume use in biological and clinical applications.^19^

An important challenge for adoption of microscale cell culture systems in biomedical research is compatibility with high resolution imaging. In contrast to Transwell^®^ inserts, a common commercially available platform for segregated coculture in which one cell type is cultured on a semipermeable membrane in an insert above the well plate, our device allows coculture of separated populations of cells on the same plane, therefore conferring the ability to view both populations under a microscope at the same time. The ability to view all cells is useful for monitoring the cells during the culture period (to observe confluence, morphology, and overall health of the culture). Further, imaging is a useful endpoint readout; in the present manuscript we demonstrate that our device enables coculture on glass coverslips, which can be removed from the device after the culture period and mounted on a glass slide for high magnification immunocytochemistry and single molecule fluorescence *in situ* hybridization (smFISH), important endpoints in biomedical research.^20^

Taken together, we present two injection molded microscale coculture devices specifically developed for easy use by biomedical researchers. Our devices are manufactured from polystyrene, which is traditionally used for cell culture in biology laboratories.^21,22^ We discuss key aspects of device design and manufacture, and present several use cases, including culture of limited primary testis cells from surgical patients and coculture to study interactions between adipocytes and prostate cancer cells; these use cases highlight the accessibility of our platform. Importantly, we have manufactured several thousand devices and sent them to eight independent biology ‘test labs,’ where they are used for wide-ranging applications including lymph, prostate, and microbial signaling. The work presented in this manuscript was collected in three independent laboratories—with devices and protocols shipped from the University of Washington to the Jorgensen lab in Comparative Biosciences at the University of Wisconsin-Madison (MA-10 cell culture and smFISH imaging) and the Crawford lab in Surgery at NorthShore HealthSystem, University of Chicago (prostate cancer-adipocyte signaling), demonstrating the robustness and accessibility of our culture platform.

## Materials and Methods

### Device fabrication

We have previously described details of design, fabrication of 3D printed and milled devices, and cleaning procedures.^9^ Briefly, devices were fabricated with a 3D printer (Form 2, Formlabs) or a CNC mill (PCNC 770, Tormach; Datron Neo, Datron) during the iterative design process. The final devices that are featured in this work were injection molded through Protolabs (Protolabs, Maple Plain, MN). The data shown in Figure 6b,e was generated with a version of the Monorail2 device that was fabricated with a CNC mill. Original design files are included in the ESI.

### Device preparation

Prior to use, all devices were rinsed with deionized water, sonicated for one hour in isopropanol and for 30 min in 70% ethanol, air dried and treated with oxygen plasma at 0.25 mbar and 70 W for 5 min in a Zepto LC PC Plasma Treater (Diener Electronic GmbH, Ebhausen, Germany). For the evaporation assays (Figures 3 and 4), devices were placed directly into the wells of tissue culture treated 12-well plates (Corning 3513). For immunostaining experiments (Figure 6), glass coverslips were sterilized with 70% ethanol, inserted into wells, and submerged in a coating solution containing 0.01 wt% poly-L-lysine (Sigma, P4707) for 30 min. Coating solution was then aspirated, and coverslips were washed 3 times with sterile deionized water. Monorail1 devices (Figure 6c,d) or milled Monorail2 devices (Figure 6b,e) were then placed on top of glass coverslips inside of wells. Collagen I was used to make the hydrogel wall for experiments shown in Figure 3 and Figure 6b,c,e. For all other experiments, low gelling temperature agarose was used to make the hydrogel wall; we recommend using agarose for experiments involving primary cells isolated from tissues digested with collagenase (Figure 5), as residual collagenase can digest the hydrogel wall if collagen I is used. Detailed protocols for preparation/use of the devices presented in this manuscript can be found in the SI. These protocols include “test your hands” sections that we send out to biology labs who use these devices. We recommend that new users follow the “test your hands” protocol (which uses food coloring added to the cell culture chambers) to ensure that they can form the hydrogel walls correctly before running biological experiments.

### Hydrogel preparation

*Collagen* was prepared using a 1:9 solution of 10X HEPES (500 mM HEPES with 10X PBS and pH 7.6): ~9 mg/mL rat tail collagen I (Corning, 354249), producing a final concentration of 1X HEPES and ~8 mg/mL collagen. The collagen solution was pipetted into the devices at the loading port. After conclusion of flow in devices, plates were incubated at 37 °C for at least 30 min before 1X PBS loading.

*Agarose* was prepared using low gelling temperature agarose (Sigma-Aldrich, 39346-81-1) and 1X PBS for a final concentration of 1.5 wt%. Gel solution was autoclaved for sterilization and to aid in dissolving agarose. Gel solution was heated to 55 °C before loading into device and allowed to cool at room temperature after conclusion of flow. Once gelled, devices were loaded with 1X PBS.

### Cell culture

#### Cell culture for imaging and viability (Figures 3, 4, and 6)

Human lung microvascular endothelial cells (HLMVEC) (Cell Applications, 540-05a) were cultured in EGM^TM^-2 endothelial cell growth media (Lonza, CC3162). MA-10 cells (ATCC, CRL-3050)^24^ were cultured in DMEM/F12 (Gibco, 11330-032) media containing 5% horse serum (Gibco,16050), 2.5% fetal bovine serum (FBS) (HyClone, SH3039603), 1.5 g/L sodium bicarbonate. BHPrS1 cells (benign human prostate stromal cells, from Simon Hayward’s lab at NorthShore HealthSystems) were cultured in RPMI-1640 medium (Gibco, 22400-089) with 5% FBS (HyClone, SH3039603).^23^ In addition, both MA-10 and BHPrS1 cell culture media were also supplied with penicillin (100 units/ml)/streptomycin (100 μg/ml) (Gibco, 15140122). Cells were cultured at 37 °C under 5% CO_2_. Cells were trypsinized (Gibco, 12604021), resuspended at 3.8 × 10^5^ cells/mL and seeded into devices at cell seeding density of 250-300 cells/mm^2^. For 6-well plate experiments (Figure 6a,e), 200 µL of sterile water was added to the edges of 6-wells containing devices; care was taken to prevent this added water from reaching the device in the center of the well. 6-well plates were placed in a bioassay dish (245 mm x 245 mm) containing about 50 mL of sterile water and incubated. For 12-well plate experiments using the Monorail1 device, 1 mL of sterile water was added in the interwell spaces (Figure 3a), and 2 mL of sterile water was added to any wells that did not contain a device. 11 µL of media was loaded into each culture chamber in the device, followed by an addition of 11 µL of cell suspension. Cell culture media was changed partially (approximately half of the media was exchanged for fresh media in each chamber) each day, and external water for evaporation control was replenished when a reduction in volume was visible. For 12-well plate experiments using the Monorail2 device, 8 and 20 µL of cell suspension were loaded into the center and outer chambers of the device, respectively, followed by 500 µL of media to the media reservoir (Figure 4aii). Cells were fixed with 4% paraformaldehyde (Fisher Scientific, AA433689M) prior to staining.

#### Primary cell isolation and culture (Figure 5)

All experiments were conducted under an institutional review board approved protocol from subjects who provided informed consent for use of residual tissue from testicular sperm extraction. Following isolation of sperm for cryopreservation, residual testis tissue was received and placed in DMEM/F12 media containing 10 mg/mL type IV collagenase and 25 mg/mL DNAse for 30 minutes at 37 °C.^25^ Following enzymatic digestion, tubule tissue settled for 5 minutes with gravity and the supernatant was removed and centrifuged for 7 minutes at 250 RPM. The resulting pellet was resuspended in culture media (Adv DMEM/F12 media with 1% BSA, 10% fetal bovine serum and 1% Pen/strep) and underwent two washings and centrifugations prior to resuspension in 100 μL of culture media. Viability of mixed interstitial testis cells was assessed with a trypan blue assay and was greater than 80% prior to seeding into the device. 8 μL of cell suspension was added to each center well of the Monorail2 device, with media placed in the outer wells and media reservoir. Primary testis cells were incubated at 34 °C for 4 and 7 days prior to fixation with paraformaldehyde for phase contrast imaging.

#### Prostate cancer cell-adipocyte coculture (Figure 7)

In this study, the experiments were performed using human prostate cancer cells (PC-3) and an adipocyte phenotype cell line (3T3-L1). The cell lines were purchased from ATCC (Manassas, VA, USA). Cells were maintained in DMEM medium (Gibco, Ref #:11330-032) containing 10% FBS and 1% penicillin/streptomycin (Gibco, 15240) in flasks. PC-3 and 3T3-L1 cells were cocultured in separate wells of the Monorail1 device. We prepared a cell density of 1 x10^6^ cells/mL and 10 μL of the solution was placed in each well of the Monorail1 device along with 12 μL of medium. The cell containing device was then incubated at 37 °C under 5% CO_2_ in RPMI medium (ThermoFisher Scientific, Waltham, MA, USA) containing 10% fetal bovine serum (FBS; Sigma, St Louis, MO, USA) and penicillin (100 units/ml)/streptomycin (100 μg/ml). After 24 hours, cells were fixed in 10% formalin prior to staining.

### Proliferation and viability Assays

For the proliferation assay shown in Figure 6c, 5-Ethynyl-2′-deoxyuridine (EdU, Invitrogen) was prepared according to the manufacturer’s specifications and diluted in cell culture media to 10 μM. EdU was added to cell culture and incubated for 6 hours; then the cells were fixed with 4% paraformaldehyde. Cells treated with EdU were subjected to Click-iT reaction cocktail (Invitrogen), which was prepared according to the manufacturer’s specifications and incubated with cells for 30 min. For assessment of cell viability shown in Figures 3 and 4, live/dead staining was performed by incubation of cells with 2 nM ethidium bromide (dead) and 10 μM Calcein AM (LIVE/DEAD^®^ Viability/Cytotoxicity Kit for mammalian cells, Invitrogen, L3224) for 30 minutes at 37 °C.

### Immunocytochemistry

For fluorescence images shown in Figure 6, cells were fixed, then devices were carefully removed from coverslips using forceps. Fixed MA-10 cells (Figure 6b), HLMVECs (Figure 6c), and BHPrS1 cells (Figure 6d) on coverslips were permeabilized with 0.5% Triton X-100 for 30 min and blocked with 3% BSA, then incubated with 2 μg/mL anti-α-tubulin antibody raised in rat (Invitrogen, MA180017) overnight at 4º C. After washing 3-5 times with PBS containing 0.1% Triton X-100, goat anti-rat secondary antibody conjugated with Alexa Fluor 488 (Jackson ImmunoResearch, 112545167, 1.5 mg/mL) for MA-10 and BHPrS1 cells, or with Alexa Fluor 647 (Jackson ImmunoResearch, 112605167, 1.25 mg/mL) for HLMVECs, was added at a 1:200 dilution and incubated with cells for 1 h followed by a 20 min incubation with 5 μg/mL Hoechst 33342 (Invitrogen, H1399). HLMVECs were stained for actin with phalloidin conjugated with Alexa Fluor 488 (ThermoFisher Scientific, A12379). Cells were washed as above. Coverslips were then placed on glass slides with VectaShield antifade mounting media (Vector Laboratories, H1000) and sealed with nail polish (Electron Microscopy Sciences, 72180). Fluorescence images were acquired using an Axiovert 200 Zeiss microscope equipped with Axiocam 503 mono camera. Phase contrast images were taken with Zeiss Primovert inverted microscope with a MU1403B camera (AmScope).

### Single molecule fluorescence in situ hybridization (smFISH)

The MA-10 cells shown in Figure 6e were washed in PBS and fixed with 4% formaldehyde for 15 min followed by permeabilization with 70% ethanol for 1 hr. After washing the cells with wash buffer for 5 min (2x SSC and 10% formamide), 50 μl of hybridization solution containing the RNA probes was added. The RNA probe sets for Star and *Cyp11A1* were generated with the Stellaris probe designer and the probes were dissolved in TE buffer, pH 8.0 (LGC Biosearch Technologies). A clean coverslip was placed over the sample to prevent drying of the hybridization solution during the incubation. The hybridization solution contained 10% dextran sulfate (Sigma, D8906), 10% deionized formamide (Ambion, AM9342) and 2x SSC (Ambion, AM9765). Samples were incubated in a dark humidified chamber at 37 °C overnight. After a 30 min wash in wash buffer, samples were incubated for 30 min in DAPI (wash buffer with 5 ng/ml DAPI) to counterstain the nuclei. After a brief incubation with 2x SSC for 5 min, antifade GLOX buffer (2x SSC, 10% glucose and 1M Tris, pH 8.0) was added without enzymes for equilibration followed by incubation with added glucose oxidase (Sigma, G2133) and catalase (Sigma, C3515) for 5 min. The samples were mounted with a drop of Prolong Gold antifade reagent (Invitrogen, P36930). Super-resolution imaging was performed with a Nikon-Structured Illumination Microscopy (N-SIM) system equipped with a SR Apo TIRF 100x objective and an iXon3 camera (Andor Technology). The images were acquired as 3D-SIM Z-stacks and analyzed using NIS-Elements software (Nikon).

### Oil-Red-O stain

PC-3 and 3T3-L1 cells were grown in the Monorail1 device on glass coverslips coated with poly-L-lysine (Sigma, St. Louis, MO, USA) and cultured and treated as described above. Cells were then washed three times with PBS, fixed in 10% formalin (30 min at room temperature), and stained with Oil-Red-O (Oil-Red-O Stain, propylene glycol; Newcomer Supply, Middleton, WI, USA) to visualize neutral lipids. Cells were also counterstained with hematoxylin (Newcomer Supply, Part # 1180G) for 10 minutes and lithium carbonate (Sigma, L4283-100G) for ~5 seconds to add contrast and highlight the nucleus. Coverslips were mounted on glass slides and sealed with Permaslip Mounting Medium (Alban Scientific Inc.). Pictures were taken of representative fields for each treatment using a 100x objective to highlight intracellular lipid droplets.

### Lipid droplet area

In slides stained with Oil-Red-O, positive intracytoplasmic lipid droplets were evaluated in 15 high power fields/experimental group. Area was calculated using Image J software (NIH, Bethesda, MD).

### Statistical analysis

To determine differences between experimental groups, a Student’s t-test was used and findings were considered significant when P<0.05. Graphs were made using GraphPad Prism, version 7.03.

## Results and Discussion

### Device overview: segregated coculture on a well plate surface

In previous work, we introduced a 3D printed platform that uses hydrogel walls patterned on a plastic or glass surface to make unique segregated coculture systems integrated into standard cell culture well plates for mammalian cell culture; we also demonstrated proof-of-concept soluble factor exchange through hydrogel using an established microbial coculture system.^9^ The present manuscript focuses on essential developments to translate this platform to biomedical laboratories, including engineering design modifications that enable manufacturing by injection molding, features for preventing evaporation, and the development of workflows that enable high resolution imaging at the end of culture. We designed two hydrogel patterning devices for culture of multiple cell types (referred to as the “Monorail1 device” and the “Monorail2 device”). We used 3D printing and computer numerical control (CNC) milling to prototype the platforms and manufactured the final devices with rapid injection molding using Protolabs. As shown in Figure 1, the Monorail1 device fits securely into the well of a 12-well plate and enables segregated coculture of up to three cell types. Pressure struts are features of the device that apply pressure to the walls of the well, allowing the well plate to be handled or inverted without dislodging the device. As shown in Figure 1, three cell culture chambers are delineated by hydrogel walls that encompass the perimeter of the cell culture chambers. When a device is placed in the well of a 12-well plate, the foot of the device (which runs around the perimeter of the device) holds the rails 250 μm above the floor of the well. A hydrogel precursor solution is loaded into the loading port of the device and flows spontaneously in the 250 µm gap between the rails of the device and the floor of the well plate. Hydrogel precursor solution flows and is confined to the space under the rail. As in our prior work,^9^ we used rails with a trapezoidal cross-section to induce pinning of the fluid on the edges of the bottom of the rail. This geometry prevents the hydrogel precursor solution from wetting the vertical faces of the device. After completion of flow, the hydrogel solution can be polymerized to yield a selectively permeable barrier that demarcates a set of chambers in which cells can be cultured (Figure 1). Cells seeded on either side of the hydrogel wall are physically separated, as shown schematically in Figure 1d, while soluble factors (e.g., small molecules, proteins) diffuse through the hydrogel wall. Up to three distinct cell populations can be seeded into different cell culture chambers. This platform can be used with several hydrogels, including collagen I and matrigel.^9^ In the present work, we developed protocols for use of low gelling temperature agarose, recommended instead of collagen when working with primary cells isolated from tissues digested with collagenase because residual collagenase can degrade the collagen wall.

**Figure 1.**
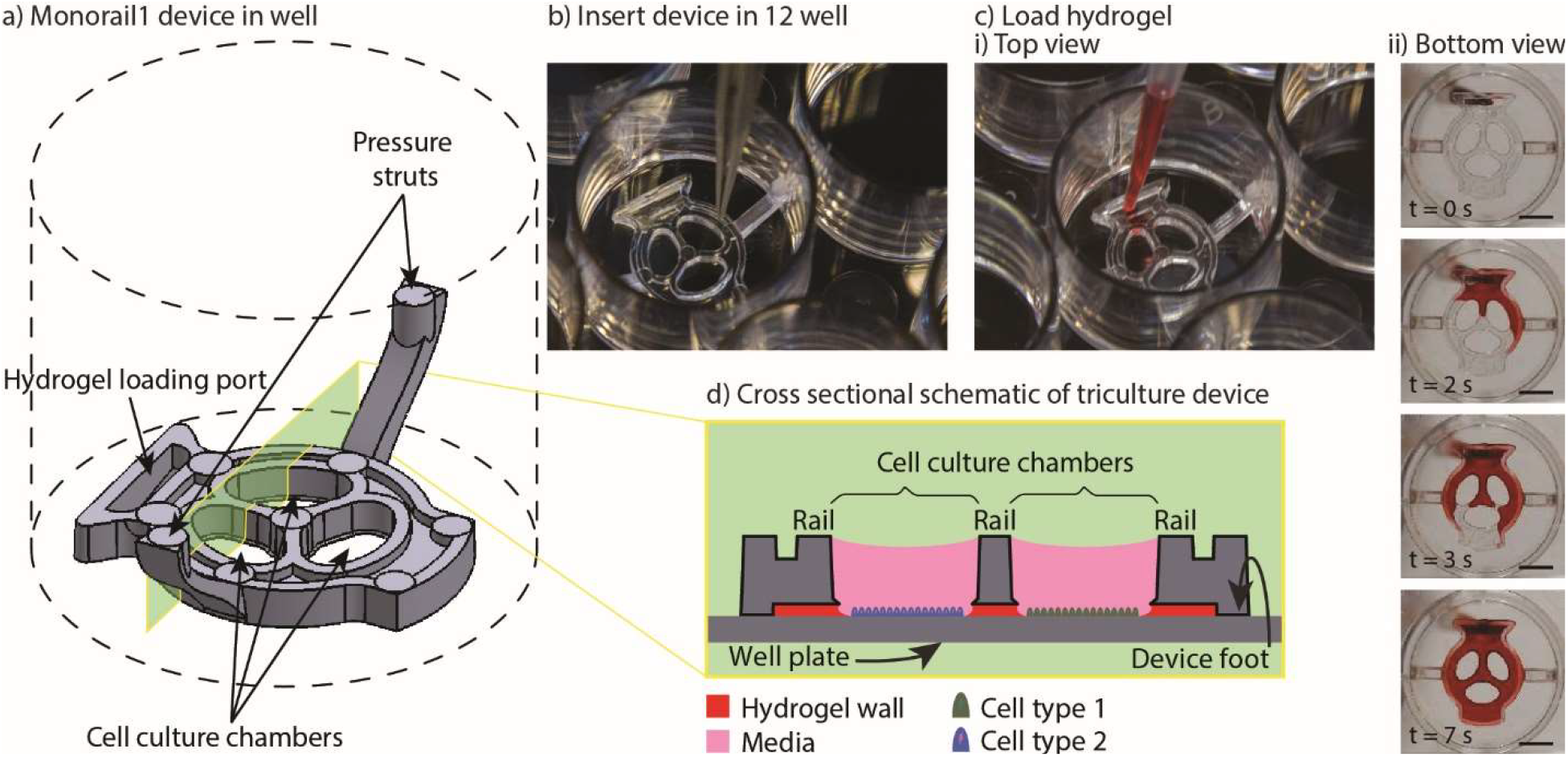
Overview of Monorail1 device design features and operation. a) Monorail1 device (dashed lines represent the well of a 12-well cell culture plate. b) The device is loaded into the well of a standard 12-well cell culture plate using forceps. c) Hydrogel is loaded into the hydrogel loading port with a standard pipet, as seen from i) above the well plate or ii) below the well plate. Hydrogel is tinted with red dye for visualization purposes. ii) Bottom view images of the gel-precursor solution patterning progression at four timepoints (video is included in the SI). Scale bars: 5 mm d) Schematic of the device in cross-section depicting distinct cell types (blue, green) in different culture chambers that are separated by a hydrogel wall formed between the bottom of the rail and the base of the well.

### Design considerations for injection molding open microfluidic devices

Our open microfluidic cell culture devices were designed for manufacture with injection molding. Advantages of injection molding include high fidelity, reproducibility, and production of large numbers of devices at a relatively low cost per device. Injection molding is a fabrication method in which the geometry of a plastic device is cut as the negative space inside a metal mold. Molten thermoplastic is injected into the negative space of the mold through an opening called a gate. Once the plastic cools, the mold is separated, the plastic device is ejected, and the mold is reused to make more devices. The ability to produce many devices from a master template (the mold) in an automated fashion makes injection molding an attractive fabrication method for high volume production of plastic parts. “Rapid” injection molding is a relatively new type of injection molding that is much cheaper than classic injection molding, making it accessible to a wide consumer-base, including academic labs. Rapid injection molding companies use computerized workflows and less robust molds (often made from aluminum) to reduce mold costs (typically ~$2,000-4,000 per mold vs. ~$50,000 or more per mold in traditional injection molding). Rapid injection molding companies typically guarantee the molds for fewer parts (~10,000 parts) in comparison to traditional injection molding which allows for high volumes (often >1 million parts). Rapid injection molding fills a specific niche in microfluidic technology development—once a design has been validated via lower volume fabrication methods such as micromilling or 3D printing, rapid injection molding can be used to translate the technology to biomedical labs that require hundreds to thousands of devices. We have previously presented a general discussion of key features important in rapid injection molding for microfluidic devices.^19^ Here we discuss aspects specific to the present device design, such as drafting, placement of ejector pins, and cored regions.

Rapid injection molding companies impose relatively stringent design constraints on parts in order to keep the mold simple and the price low. The most important of these constraints is that every face of the device must be visible from either the top or the bottom of the part. This necessitates every vertical face of a device to be drafted (i.e., angled slightly); see computer aided design (CAD) files and schematics included in the SI. When this criterion is met, the device can be fabricated with a two-sided mold. Importantly, because our devices use open microfluidics rather than closed channels, the entire device is fabricated as a single injection molded part without subsequent bonding steps. Figure 2 shows the surfaces of each device that are defined by either the “A” side (red) or the “B” side (green) of the mold. When the A and B sides of the mold come together, the space in between them is the void that the molten plastic fills, which ultimately becomes the molded plastic device (Figure 2). Two-sided molds offer the simplest and cheapest incarnation of rapid injection molding. There are more complicated incarnations of rapid injection molding that are less stringent on device design but more expensive. Figure 2 also shows the locations of gates, where molten plastic is forced into the mold during the injection molding process. For the devices presented here, gates are located on the backs of the devices so that any defects from post processing do not affect device performance.

**Figure 2.**
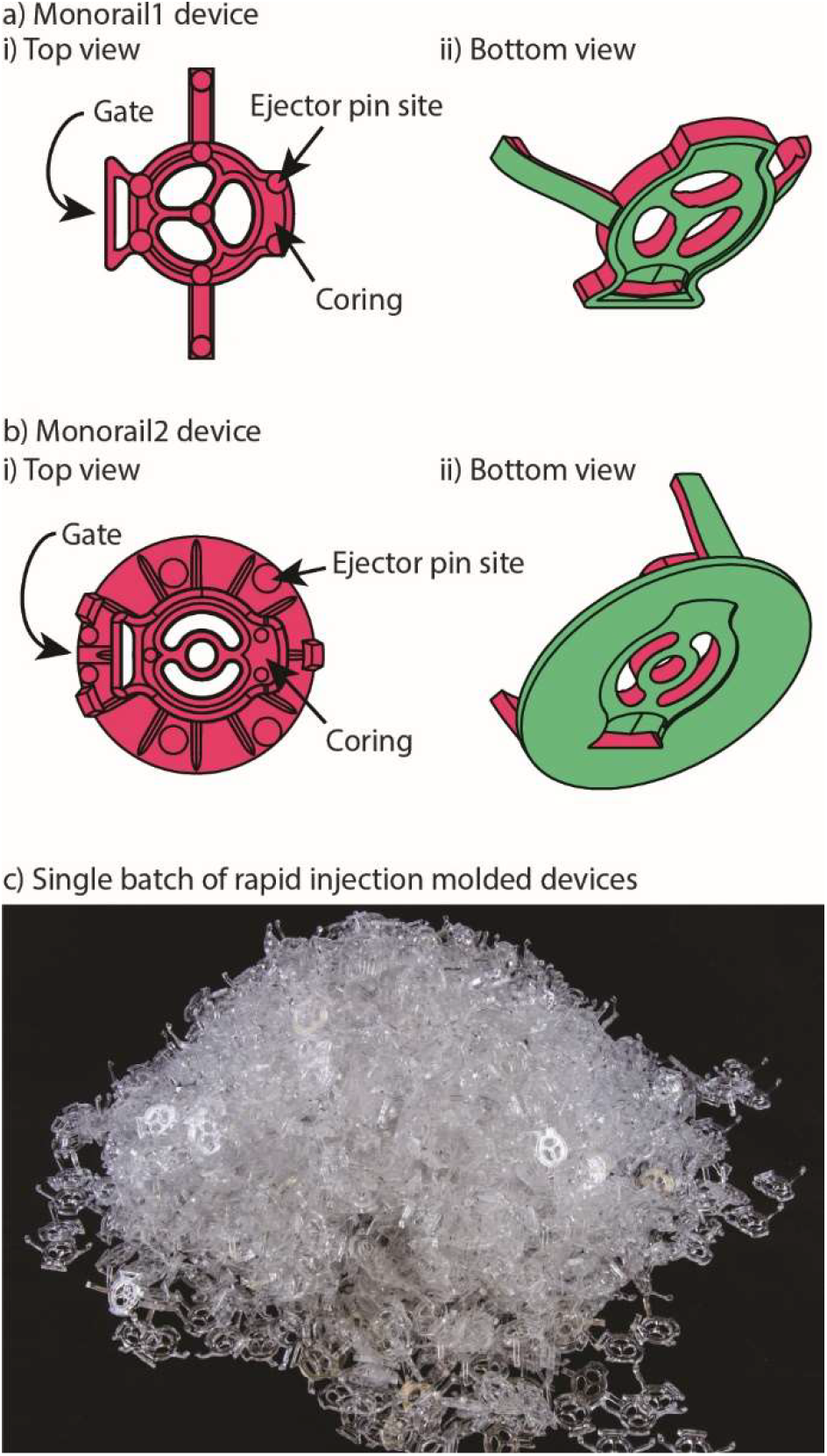
Monorail devices can be fabricated via rapid injection molding. a) Monorail1 device and b) Monorail2 device schematics showing location of coring, ejector pins (all small round circles), and gates from top (i) and bottom (ii) views. c) Photo of Monorail1 devices fabricated with rapid injection molding.

After the device is molded, the molten plastic is allowed to cool. The cooling process can cause shrinking in thicker areas of the device due to differential cooling time between thick and thin areas in the device. Shrinking manifests as sunken areas in a device, where surfaces of the device that were designed to be flat come out as concave and sunken into the device. Coring out thick areas of the device creates more uniformity in the thickness of the device, mitigating shrinkage anomalies (examples of cored regions are indicated in Figures 2ai and 2bi). Once the device is cooled and the two sides of the mold are separated, the device must be ejected from the mold. This is accomplished with ejector pins, which push the device out of the mold after the molding process is complete. The placement of ejector pins in the mold is important so that an even force is applied across the entire device. Devices that are fabricated with injection molding must be designed with space for these ejector pins to push against, which manifest as small circles in the final device. This informed our decision to design the ejector pins to be located on the top (“A” side) of the device so that they would not interfere with the open microfluidic hydrogel patterning that occurs on the bottom (“B” side) (Figure 2a).

### External evaporation control: adding water within the well plate

Open microfluidic systems offer the advantage of total pipet accessibility while closed systems are accessible only from strategically placed ports. However, the relatively large area of exposed liquid surface makes open microfluidic systems more susceptible to the deleterious effects of evaporation.^13,14^ In most cases, researchers circumvent this problem by incubating their microculture systems in a petri dish with water droplets surrounding the device and then placing the petri dish in a secondary container with additional water (typically 5-100 mL) which keeps the partial pressure of water vapor in the containment unit near equilibrium, thus mitigating evaporative water loss in the culture system.^6,26^ Other secondary containment strategies involve surrounding the microculture-ware with wetted Kimwipes^TM^ in a larger container. A disadvantage of this approach is that the experiment can become contaminated in multi-day culture experiments. While the microscale cell culture system itself may be quite small, a large secondary containment system often uses a lot of space in the cell culture incubator and comes with an inherent risk of liquid spilling. Finally, the need for secondary containment can be cumbersome in the hands of researchers that are not accustomed to microfluidic devices. To address these disadvantages, we optimized a protocol to mitigate evaporative loss in the Monorail1 device without the need for secondary containment. We cultured testis cells (MA-10) in the device and used cell viability as a qualitative metric to assess evaporation. Negligible evaporation was inferred in the setting of high cell viability. We found that cells were nearly 100% viable after 24 hours in culture when the four corner wells, as well as the spaces between the wells, were used as reservoirs for additional water, as shown in Figure 3a. If the user loads fewer than eight devices into a well plate, we recommend adding additional water to vacant wells. The eight-device layout maximizes the number of usable wells in a well plate while keeping cell viability high. Other layouts showed dispersed pockets of dead cells throughout the well plate, likely due to evaporation in the microculture system (Figure S3). Multiple cell types were successfully cultured and showed comparable viability in the Monorail1 device (Figure 3c-d). Without the need for secondary containment to address evaporation concerns, this device offers a simple platform for coculture and triculture experiments in a form factor that is familiar to biomedical researchers.

**Figure 3.**
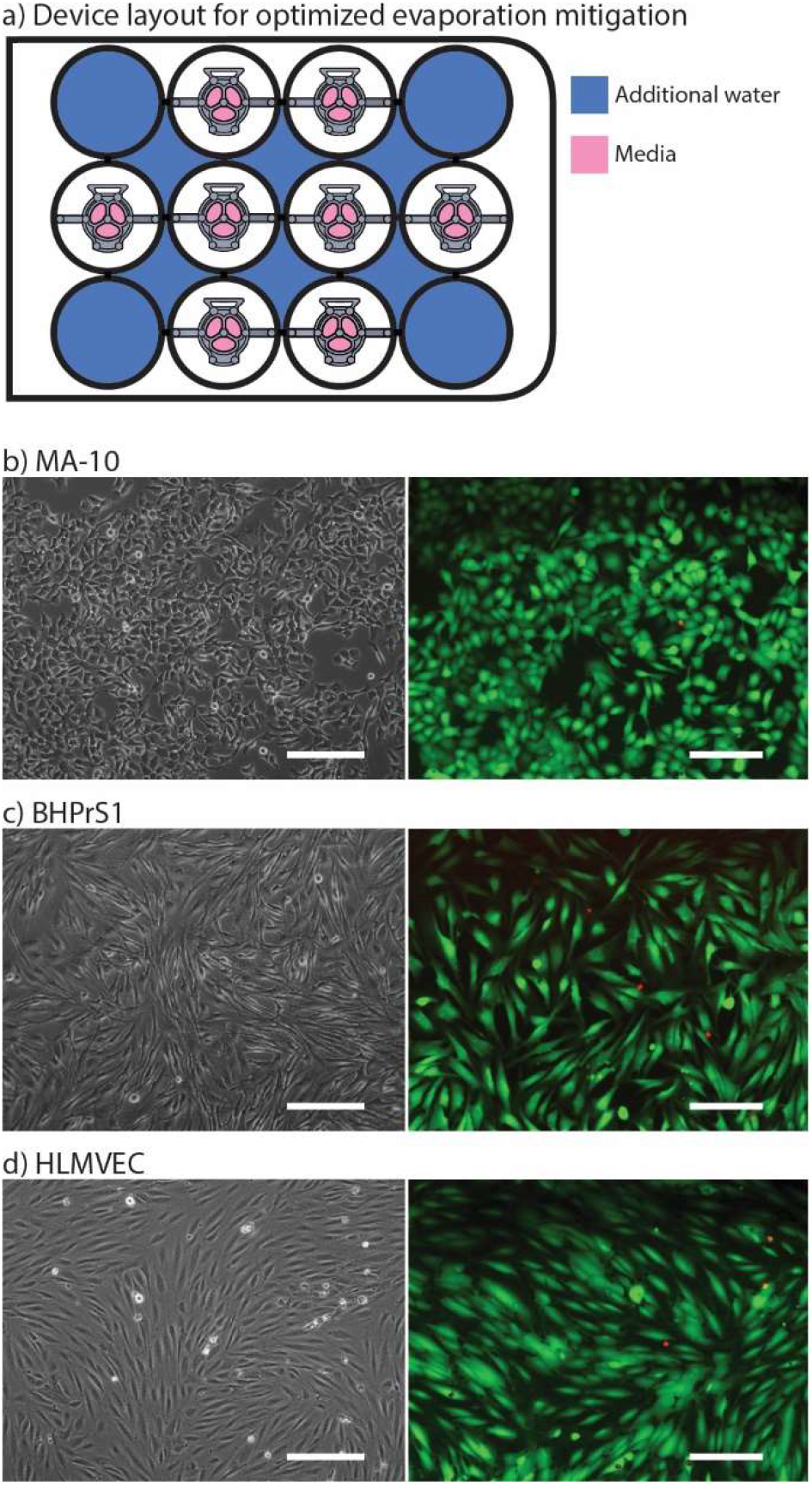
Adding water to corner wells and interwell spaces mitigates evaporation. a) Schematic of Monorail1 device layout in 12-well plate that optimizes evaporation control. Blue color indicates well or interwell space that is filled with water to mitigate evaporation. b) Testis cells (MA-10), c) benign human prostate stromal cells (BHPrS1), and d) primary human lung microvascular endothelial cells (HLMVEC) were cultured in the Monorail1 device for 24 h in the layout shown in a. Left images are phase contrast microscopy showing expected cell morphology. Right images show results of live (green) and dead (red) cell staining performed at 24 h. Fluorescence images are representative of the lowest viability field of view that was observed. Scale bars: 200 µm.

### Internal evaporation control: within-well evaporation control

We also developed a microscale coculture device with a smaller cell culture area and in-device evaporation control. This second device (referred to as the Monorail2 device) was conceived to enable coculture experiments with rare cells, or cultures of cells that require soluble factors from supporting cells to maintain viability *in vitro* (Figure 4). To address this need, the Monorail2 device features a smaller central cell culture region (~3 mm^2^) flanked by two larger outer culture regions (~8 mm^2^ each), which hold 8 and 20 µL of media, respectively. Figure 4 shows details of the Monorail2 device, which was designed with a built-in media reservoir. The media reservoir surrounds the periphery of the cell culture chambers and is segregated by a pinning ridge. When the device is secured in the bottom of the well, there is a thin void space under the device, created by imperfections in the plastic surfaces of the device and well (labelled “contact area” in Figure 4a ii). When fluid evaporates from the cell culture chambers, media flows under the device from the media reservoir to the culture chambers. Importantly, we designed the device to limit diffusion of soluble factors secreted by cells in the culture chambers (or drug treatments applied in the culture chambers) to the media reservoir by making the “contact area” as large as possible within the footprint of the well. The large contact area increases the diffusion distance, mitigating diffusion on the timescale of cell culture experiments. As shown in Figure 4a ii, hydrogel precursor solution floods under the contact area when it is loaded into the device, further mitigating diffusive loss of soluble factors to the media reservoir; however, the relatively high viscosity of most hydrogels precludes them from completely filling this space. Because the concentration of soluble factors is important in biological experiments, we characterized diffusion from the cell culture chambers to the media reservoir. The diffusion of a 10 kDa fluorophore, which was used as a model soluble factor, is limited to 2.4± 0.2% after a 24 hour incubation (mean ± standard deviation for three replicates). Therefore, soluble factor diffusion from the cell culture chambers to the media reservoir is minimal, maintaining the benefit of the low volumes when pharmacologic manipulations are used in experiments.

**Figure 4.**
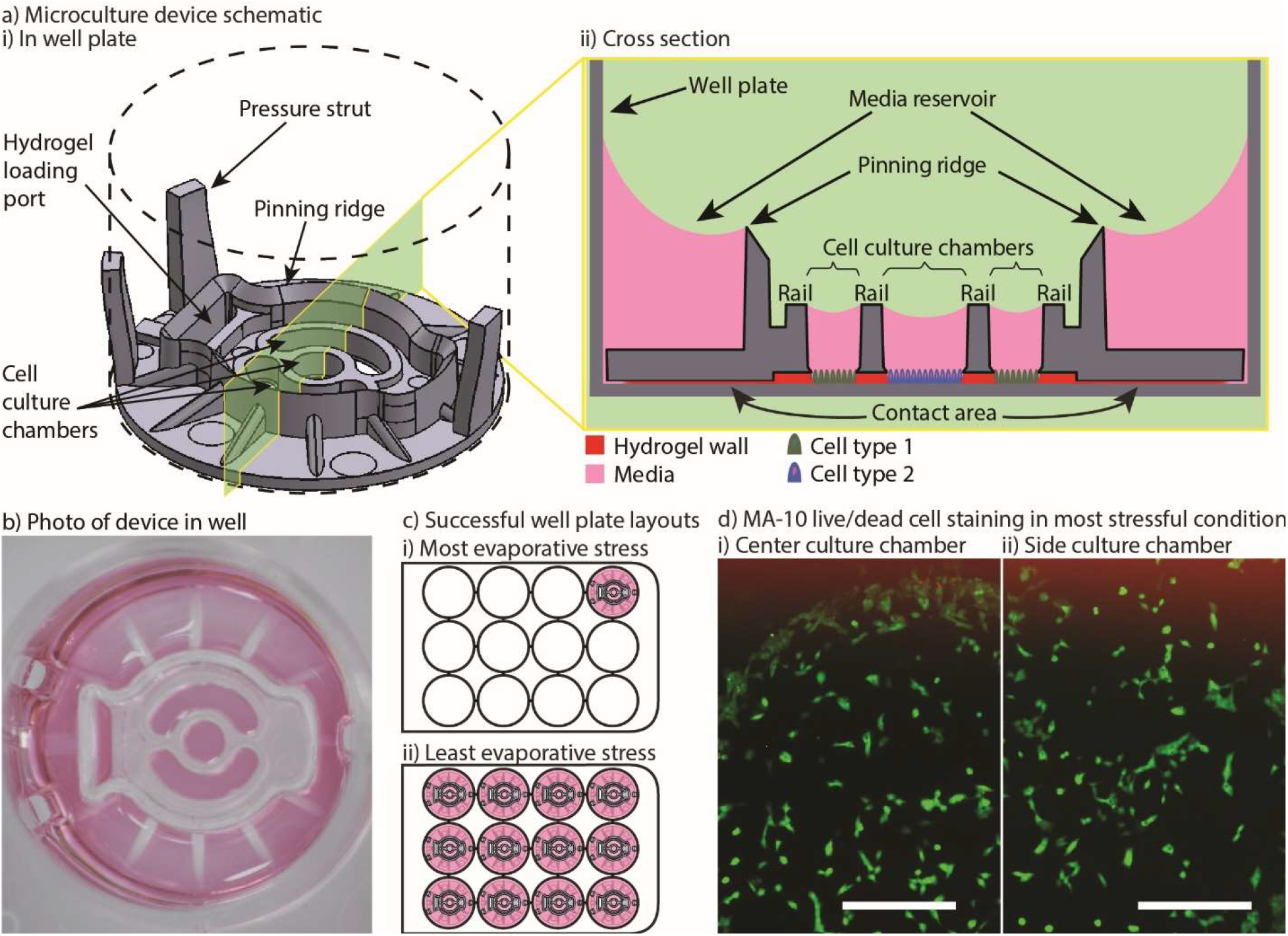
Monorail2 device design features minimize evaporative stress on cells in culture. a) i) Monorail2 device schematic (dashed lines represent the well plate of a 12-well cell culture plate). ii) Cross-sectional schematic of the Monorail2 device when loaded in a 12-well plate with media loaded into cell culture chambers. The media reservoir exists on the peripheries of the device, and is kept from spilling over the top of the device into the cell culture chambers by a pinning ridge that runs around the cell culture chambers and loading port. b) Photo of a Monorail 2 device in a 12-well plate loaded with media. Scale bar: 5 mm. c) Schematics of well plate layout experiment to test evaporation. d) Testis cells (MA-10) were cultured in the Monorail2 device for 24 hours, and live (green) and dead (red) staining was performed. High viability (nearly 100%) was observed in all device and in all configurations. Representative images of cells in the center and side cell culture chambers are shown. Scale bars: 200 µm.

To validate the mitigation of evaporative water loss in the Monorail2 device, we used the aforementioned cell viability readout and observed little to no cell death in any part of the device, irrespective of the number of devices in the well plate or the position of a device within the well plate. It is known that using a single well of a well plate offers the most challenging condition for minimizing evaporation. This is because each well of a well plate that contains evaporating water will contribute some water vapor to the other wells of the plate. Therefore, we performed cell viability experiments in well plates with devices in every well (easiest condition for evaporation control) and well plates with only one device in the corner (most challenging condition for evaporation control). As shown in Figure 4, excellent viability was observed in the most challenging evaporative condition. Of note, when using this device, it is essential to use the device as designed (i.e., placing media in the media reservoir); without media in the media reservoir, fluid will flow from the cell culture chambers to the outer ring which is incompatible with cell viability.

Some experiments utilizing primary mammalian cells require the use of enzymatic tissue digestion as a step prior to cell isolation and/or purification. We observed utilizing a collagenase digestion step could lead to digestion of the collagen hydrogel walls after 24 hours in culture; therefore, we developed a protocol for an alternative hydrogel, low gelling temperature agarose. Figure S1 shows that diffusion of molecules of 75 kDa, 10 kDa and 527 Da size is comparable with collagen or low temperature gelling agarose walls. The Monorail2 device is particularly beneficial for culture of small numbers of cells in small media volumes, or coculture of rare cells with supporting cells to study paracrine interactions. It is readily adopted by biomedical researchers because it is a well plate compatible and pipet accessible format, and has the potential to simplify a wide range of difficult coculture conditions.

We are currently working in collaboration with the University of Washington Male Fertility Laboratory to culture primary human testis cells in the Monorail2 device. Cells are collected from residual tissues after patients undergo testicular sperm aspiration (TESA) or testicular sperm extraction (TESE), surgical protocols commonly performed to collect sperm for *in vitro* fertilization or intracytoplasmic sperm injection. For these experiments, the tissues are removed in the operating room and processed in the adjacent clinical laboratory at the Male Fertility Laboratory; the ability to use our Monorail2 device in a clinical laboratory underscores the ease with which the device can be setup and operated. Here, we show preliminary results indicating that our Monorail2 device enables culture and maintenance of mixed interstitial testis cells isolated from testicular sperm extraction (Figure 5). Due to the low cell yield from this TESE, we only recovered cells sufficient to seed the central chamber of five Monorail2 devices, underscoring the importance of the microscale culture dimensions. As collagenase was used for enzymatic digestion, we used low gelling temperature agarose to make the hydrogel wall. Phase contrast images were taken of fixed cells after four and seven days in culture (Figure 5). In future work, we are developing protocols to isolate, characterize, and culture Leydig cells, a rare cell type that produces steroid hormones, including testosterone, in the testis (which will be cultured in the center chamber of the Monorail2 device) with Sertoli cells and other supporting cell types (which will be cultured in the outer chambers), paralleling our prior work with mouse fetal Leydig and Sertoli cells.^26^.

**Figure 5.**
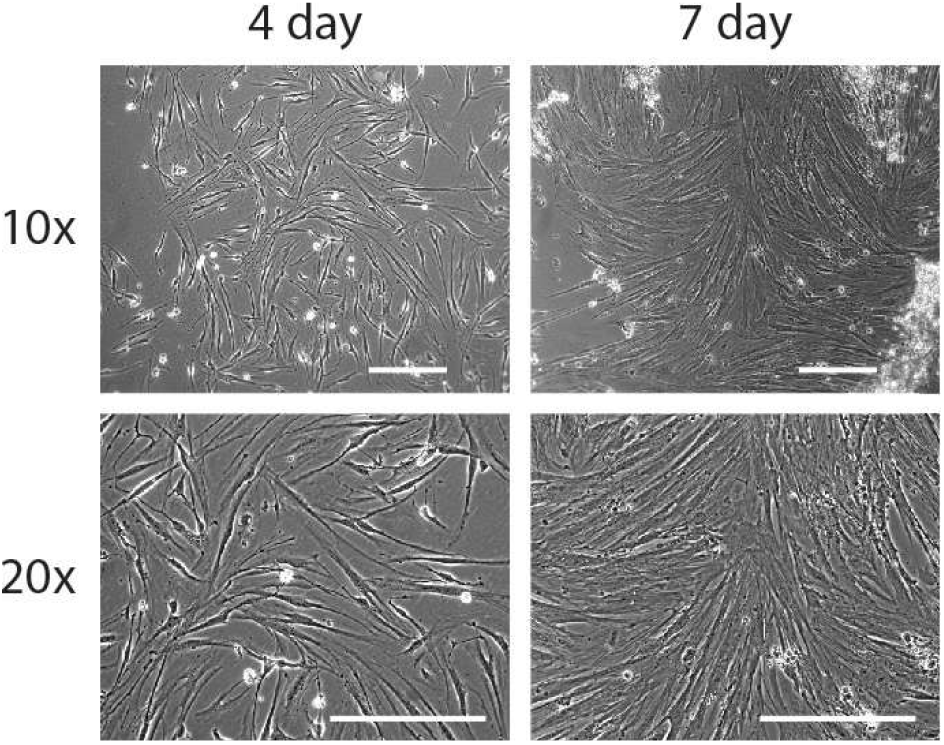
Human primary testis cells cultured in the microculture device. Cells were maintained for four days (left column) and seven days (right column), showing the ability to culture limited numbers of primary cells isolated from surgical procedures. Images are representative of three replicate devices (at day four) and two replicate devices (at day seven). Scale bars: 200 µm.

### Device removal workflow for high resolution microscopy

High resolution imaging is an important readout for biomedical researchers, as the intracellular location of the substance of interest (e.g., protein, mRNA) has important implications for function. Immunocytochemistry is a common molecular technique used to localize protein antigens of interest in cells by binding of a specific antibody. *In situ* hybridization is an analogous technique to localize nucleic acids in cells to discover both temporal and spatial information about gene expression. Single molecule fluorescence *in situ* hybridization (smFISH) detects a specific mRNA transcript with multiple short oligonucleotide probes and offers improved resolution and quantitation compared to traditional *in situ* hybridization and immunocytochemistry.^20^ It is a powerful tool for understanding the spatial and temporal patterns of gene expression at the level of the individual cell. For high resolution imaging, such as smFISH, cells are typically cultured on glass coverslips, rather than plastic cultureware since plastic cultureware limits high resolution imaging due to thickness, autofluorescence, and defects in the plastic surface. A common workflow for immunocytochemistry or smFISH sample preparation involves fixing and processing cells on a glass coverslip and then mounting the coverslip on a glass slide such that the sample is sandwiched between the coverslip and the slide. Often, samples are mounted using “mounting media”, and the coverslip is sealed to the glass slide, enabling the samples to be stored long term. Since this method is commonplace in many biology labs, many products have been developed around this workflow including holders that enable staining and chemical processing of multiple coverslips at a time, microscope stages designed to hold a standard glass slide, and boxes designed to hold glass slides that biology labs routinely use to archive samples for many years. In our prior work we showed that hydrogel patterning works on glass surfaces, such as well plates with integrated glass bottoms;^9^ here we developed procedures to use our devices on top of a glass coverslip within a well, remove the device from the coverslip after the culture period, and mount the coverslip on to a glass slide. This is also important because some specialty microscopes used for smFISH require the use of proprietary coverslips with specific thickness tolerances.

We designed devices that are compatible with two standard coverslip sizes: round coverslips (20 mm in diameter) that fit in a 12-well plate and square coverslips (22 x 22 mm) that fit in a 6-well plate. The Monorail1 and Monorail2 devices (shown in Figures 1-4) fit in a 12-well plate and are compatible with 20 mm diameter coverslips. We CNC milled an adapted device design that fits in a 6-well plate for use with 22 x 22 mm coverslips (Figure S2; design files included in the ESI). In all designs, the placement of the hydrogel loading port was an important consideration; in contrast to our prior designs,^9^ where the loading port was at the perimeter of the well, here we moved the loading port away from the edge of the well to prevent hydrogel from creeping between the coverslip and the bottom of the well plate. Figure 6a shows a schematic workflow of how samples are prepared for high resolution imaging, as well as images of cells cultured in the microscale devices. Figures 6b-d show three different cell types imaged at 63x magnification, with staining for α-tubulin (all), proliferating nuclei (Figure 6c) and actin (Figure 6d). These images show no visual distortion of immunostained cells by the hydrogel residue, supporting the compatibility of the Monorail devices with high resolution imaging.

**Figure 6.**
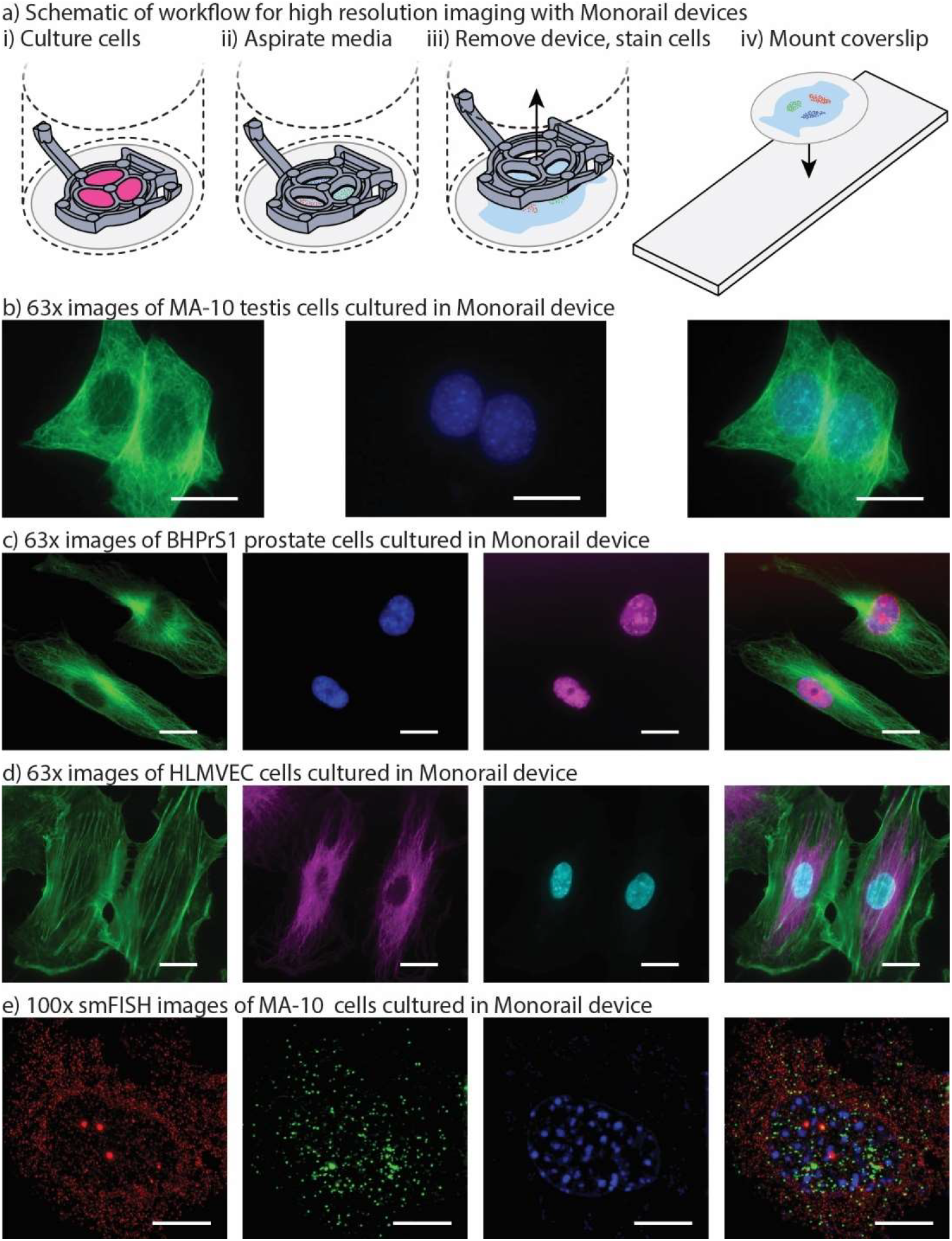
Monorail devices allow high resolution microscopy for coculture experiments. a) Schematic workflow for high resolution imaging with monorail devices. i) A glass coverslip is placed in the well of a well plate, a device is placed over the top, and cells are cultured. ii) At the conclusion of the cell culture experiment, the media is aspirated. iii) After fixing cells, the device is lifted gently from the coverslip and removed, with some hydrogel residue remaining on the coverslip; immunostaining is performed iv) The coverslip is removed from the well, inverted, and placed on a glass slide. Imaging is then performed directly through the glass coverslip by inverting the slide-coverslip assembly. This sample preparation was carried out for all high resolution images. b) MA-10 testis cells were cultured in the milled Monorail2 device in a 6-well plate and immunostaining was performed to detect α-tubulin (green) and nuclei (Hoechst, blue). Scale bars: 20 µm. c) BHPrS1 cells were cultured in the Monorail1 device in a 12-well plate and immunostaining was performed to detect α-tubulin (green), nuclei (Hoechst, blue) and proliferating nuclei (EdU, pink). Scale bars: 20 µm. d) HLMVECs were cultured in the Monorail2 device in a 12-well plate and immunostaining was performed to detect actin (green), α-tubulin (magenta), and nuclei (Hoechst, blue). Scale bars: 20 µm. e) N-SIM Z-stack (total 26 planes) images taken of smFISH probes designed to recognize Star (red) and *Cyp11a1* (green) mRNA within MA-10 cells cultured in milled Monorail2 device. Nuclei are counterstained with DAPI (blue). Scale bars: 10 µm.

Figure 6e shows single molecule fluorescent *in situ* hybridization (smFISH) performed on cultured MA-10 cells that was used to detect transcription of the genes encoding steroidogenic acute regulatory protein (*Star*) and cholesterol side-chain cleavage enzyme (*Cyp11a1*). STAR is the transport protein that translocates cholesterol from the outer mitochondrial membrane to the inner mitochondrial membrane where *Cyp11a1* is present to convert cholesterol to pregnenolone. This transport is considered the rate limiting step in steroid synthesis. The FISH probe sets contain multiple (20-40) single labelled oligonucleotides (20mers) that hybridize in series, which is necessary to facilitate single molecule resolution of transcript detection. Single mRNA transcripts are visualized as discreet spots (red or green) in the cytoplasm and are used to accurately quantify the number of transcripts for each gene. The power of this method is that it can be used to compare transcriptional responses from external signals, thereby enabling quantitative mechanistic studies.^20,27^

### Prostate cancer cell-adipocyte coculture

Obesity is known to synergize with several different types of cancer to yield higher likelihoods for tumorigenesis in cancer-free patients and enhanced tumor growth and metastasis rates in patients with cancer.^28^ As such, adipocytes have received growing attention in cancer research for their role in promoting cancer progression.^29^ In addition, inhibition of lipogenesis in prostate cancer cells has been shown to suppress tumor growth.^30^ The role of paracrine signaling between cancer cells and adipocytes has been studied, but more work needs to be done to parse out the mechanisms by which adipocyte signaling influences malignant tumors.^31^ In this work, we show preliminary data from a model of prostate cancer that is influenced by adipocyte signaling. We used the Monorail1 device to coculture prostate cancer (PC-3) cells with adipocytes (3T3-L1). Given that lipid metabolism is important to the maintenance of prostate cancer energy homeostasis, we quantified lipid droplet area in PC-3 cells in the presence and absence of 3T3-L1 cells cultured in a neighboring chamber of the Monorail1 device. Figure 7 shows an increased area per lipid droplet in PC-3 cells cocultured with 3T3-L1 cells compared monoculture, supporting the hypothesis that 3T3-L1 cells secrete soluble factors that augment lipid droplet induction in PC-3 cells.

**Figure 7.**
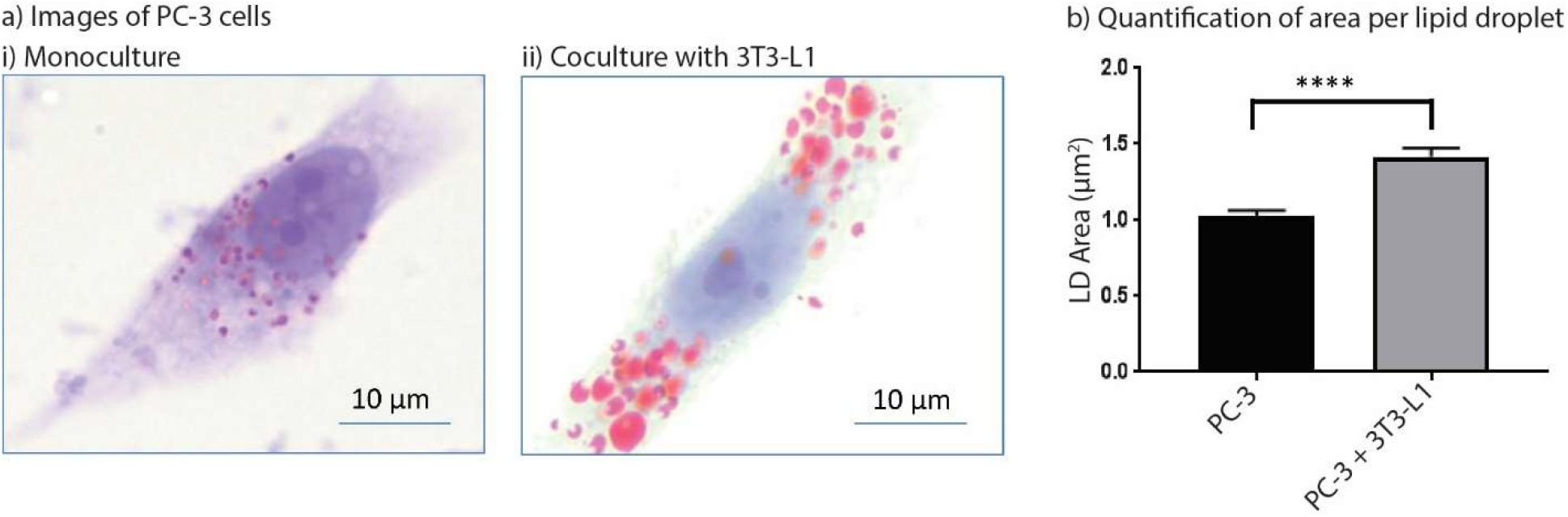
Coculture of prostate cancer cells (PC-3) with adipocytes (3T3-L1) leads to increased lipid droplet area in prostate cancer cells. a) Representative images of PC-3 cells in monoculture (i) and cocultured with 3T3-L1 (ii) (Oil-Red-O, red; counterstained with hematoxylin and lithium carbonate, blue). b) Quantification of lipid droplet area in PC-3 cells cultured alone or with 3T3-L1 cells. Lipid droplets were quantified from 15 high power (100x) fields of view for both monoculture and coculture groups. Data plotted are from one experiment and are representative of data collected across two independent experiments. Error bars represent the standard error of the mean.

## Conclusion

In this work, we developed two open microfluidic well plate inserts for coculture that are compatible with mass production via injection molding. These devices seamlessly integrate into standard well plate monoculture procedures, enabling more advanced experimentation without drastically altering experimental conditions. By manufacturing our devices with rapid injection molding, we have been able to disseminate them to numerous biology laboratories for diverse coculture applications. We find that biology laboratories with no prior microfluidics experience can readily use our devices for advanced coculture applications, including use with human primary cells coming from patients, by first following a simple protocol that they use to train and “test their hands” with food coloring (see Device Protocols included in the SI). In the future, we will continue optimizing our devices by iterating on their designs based on user feedback. Our ultimate goal is to make the platform commercially accessible, and thus add to the toolbox of cell culture technologies that are available to any biology researcher.

## Supporting information

Electronic Supporting Information

Fig 6b original image

Fig 6c original image

Fig 6d original image

Fig 6e original image

Video of hydrogel flow in Monorail1 device

Monorail1 use protocol

Monorail2 use protocol

## Acknowledgements

This work was funded by NIH grants R35GM128648 (ABT), R01HD090660 (JSJ, CRJ, ABT), R01CA242920 (OF), and R01 CA200064 (SEC), as well as a STEP Grant from the University of Washington technology transfer office, CoMotion. We thank Evan Clasen for his help with fluorescent dye diffusion experiments.

## Conflicts of Interest

The authors acknowledge the following potential conflict of interest in a company pursuing open microfluidic technologies: ABT: Stacks to the Future, LLC.

## Table of Contents Entry

An injection molded coculture platform is presented with use cases that highlight the accessibility and enabling facets of our platform.

