## Supplementary material for "Injection molded open microfluidic well plate inserts for user-friendly coculture and microscopy": Electronic Supporting Information

Table of Contents

Figure S1: Diffusion of fluorescent dyes through collagen and agarose gel walls
Figure S2: Representative images of high resolution imaging workflow
Figure S3: Representative images of sub-optimal device layout in 12-well plate
Figure S4: Monorail1 schematic showing a drafted features.
High resolution TIFF images (All)
CAD files
Protocols
Hydrogel loading video

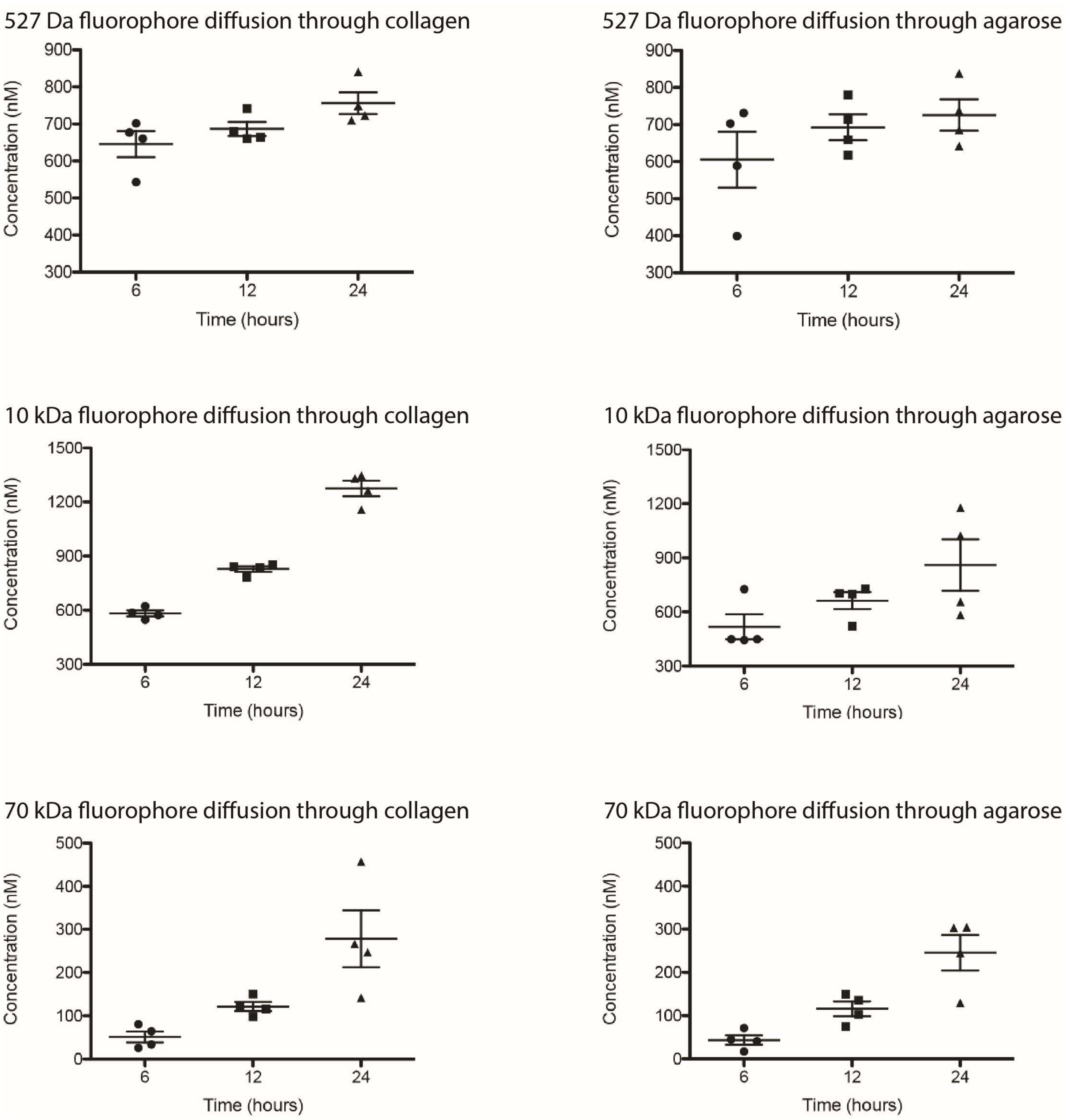


*Figure S1:* Diffusion of fluorescent dyes through collagen and agarose gel walls in Monorail2 devices. Concentrations of 527 Da (top row), 10 kDa (middle row), or 70 kDa (bottom row) fluorophore in receiving chambers of microculture device after 6, 12, and 24 hours. Gel walls through which diffusion occured were made of either type I collagen (left column) or agarose (right column). Each plotted point represents a pooled sample from both outer chambers in a Monorail2 device. Error bars represent standard deviation.

**Materials and methods for diffusion experiments (Figure S1):**

**Device preparation**

Injection molded Monorail2 devices were sonicated in isopropanol for 1 h, soaked in 70% ethanol for 30 min, and allowed to air-dry overnight. Before use, devices were plasma treated for 5 min at 0.25 mbar and 70 W in a Zepto LC PC Plasma Treater (Diener Electronic GmbH, Ebhausen, Germany) using oxygen. The devices were inserted into the bottom of a tissue culture treated polystyrene 12-well plate (Corning, 07-200-82) and loaded with 40 µL of hydrogel. Once gelled, 1X PBS was loaded into the center chamber (8 µL), side chambers (20 µL/chamber), and sacrificial media reservoir (500 µL) and stored at 6 °C overnight. 1X PBS in the center chamber was replaced with fluorescent dye the following morning (t = 0 hour), and the plate was incubated at 37 °C. Solution from the side chambers were collected and pooled at t = 6 h, 12 h, and 24 h for each technical replicate. The fluorescence of each sample was measured using a Multiskan Spectrum UV/Visible Microplate Reader (Thermo Labsystems, Waltham, MA) (n = 4 for each time point).

**Fluorescent dye preparation**

10 µM fluorescent dye solutions for Alexa Fluor 488 (MW 546 Da, Thermo Fisher, A33077), dextran Alexa Fluor 527 (MW 10 kDa, Invitrogen, D22911) and dextran fluorescein (MW 70 kDa, Invitrogen, D1823) were all prepared in 1X PBS.

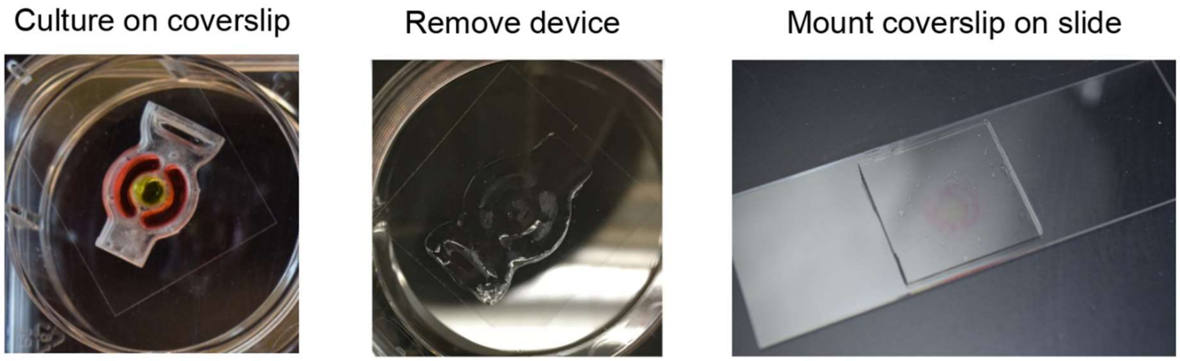


*Figure S2:* Representative photos of crucial steps in high resolution microscopy sample preparation with CNC milled Monorail2 devices designed for use in a 6-well plate. Briefly, the device is placed on a coverslip in a 6 well plate (left), and a coculture experiment is carried out (here, food dye was added to cell culture chambers for visualization). The device is then removed, leaving a faint hydrogel residue (middle). Finally, the coverslip is removed from the well with fine-tip tweezers, inverted, and placed on a glass slide (right).

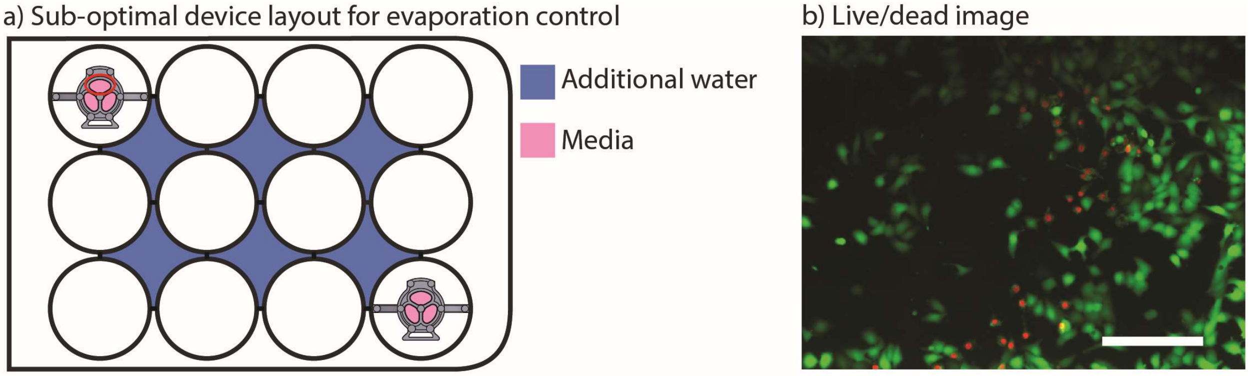


*Figure S3:* Representative cell viability in sub-optimal conditions for evaporation control. a) Well plate layout that leads to evaporation in cell culture chambers of monorail devices. b) Representative image of cell viability in top culture chamber of upper left monorail device from well plate layout.

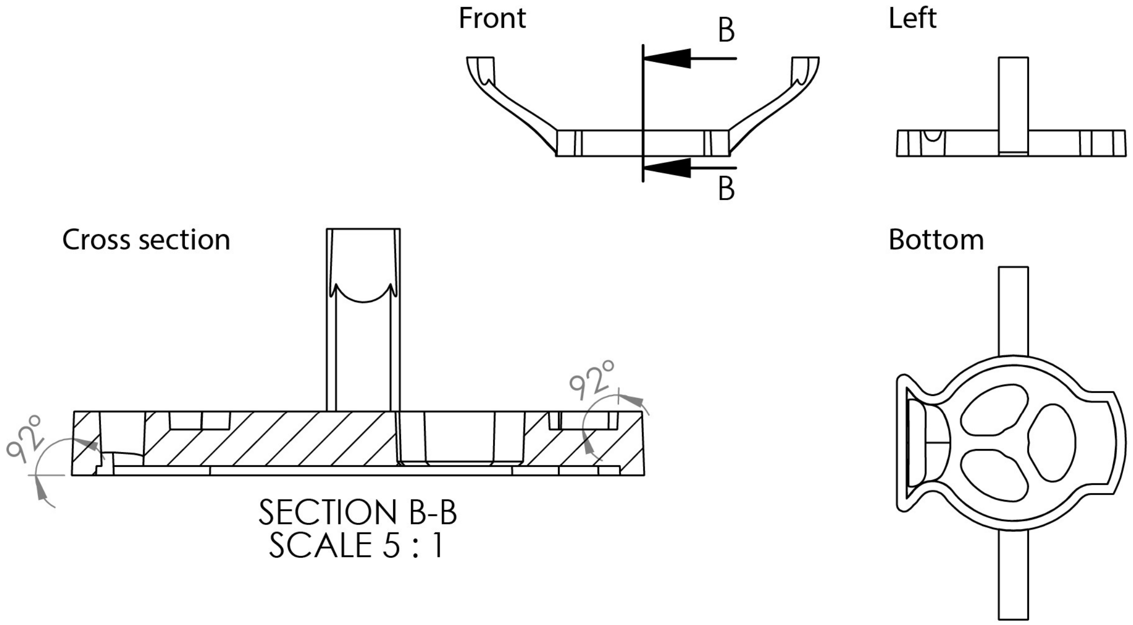


*Figure S4:* Monorail1 schematic showing a drafted features. All vertical surfaces of both injection molded Monorail devices were angles by at least two degrees to allow for demolding, the last step of the injection molding process. Cross section (left) shows two surfaces that were drafted by two degrees (92$^o$ is two degrees greater than the angle that would yield an undrafted surface).
