## Supplementary figures and images for "Injection molded open microfluidic well plate inserts for user-friendly coculture and microscopy"

### Fig 6b original image

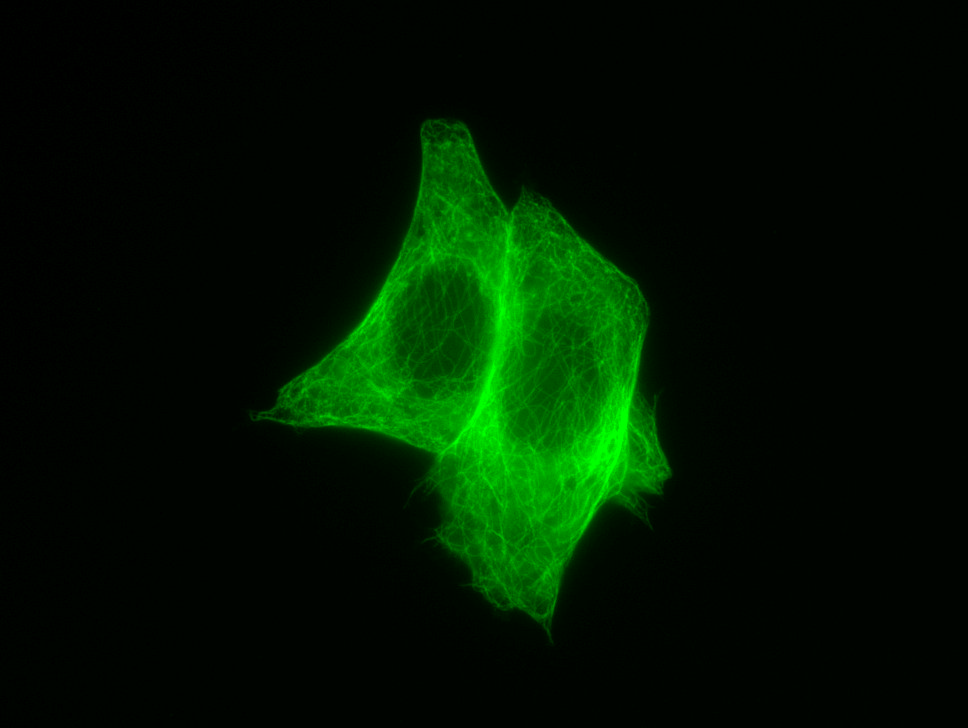

### Fig 6c original image

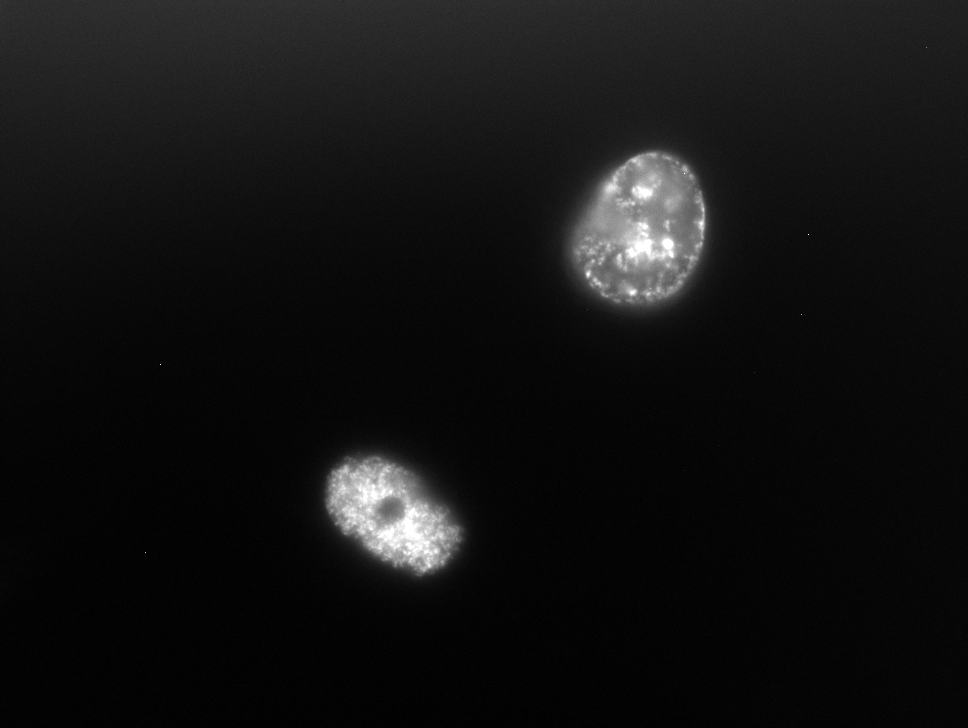

### Fig 6d original image

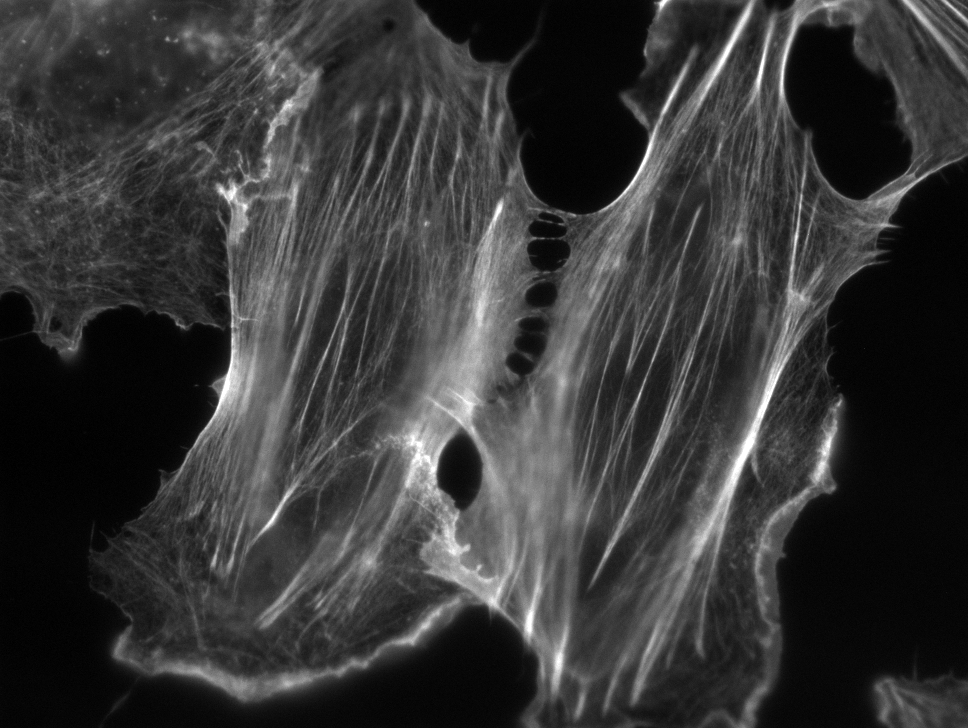

### Fig 6e original image

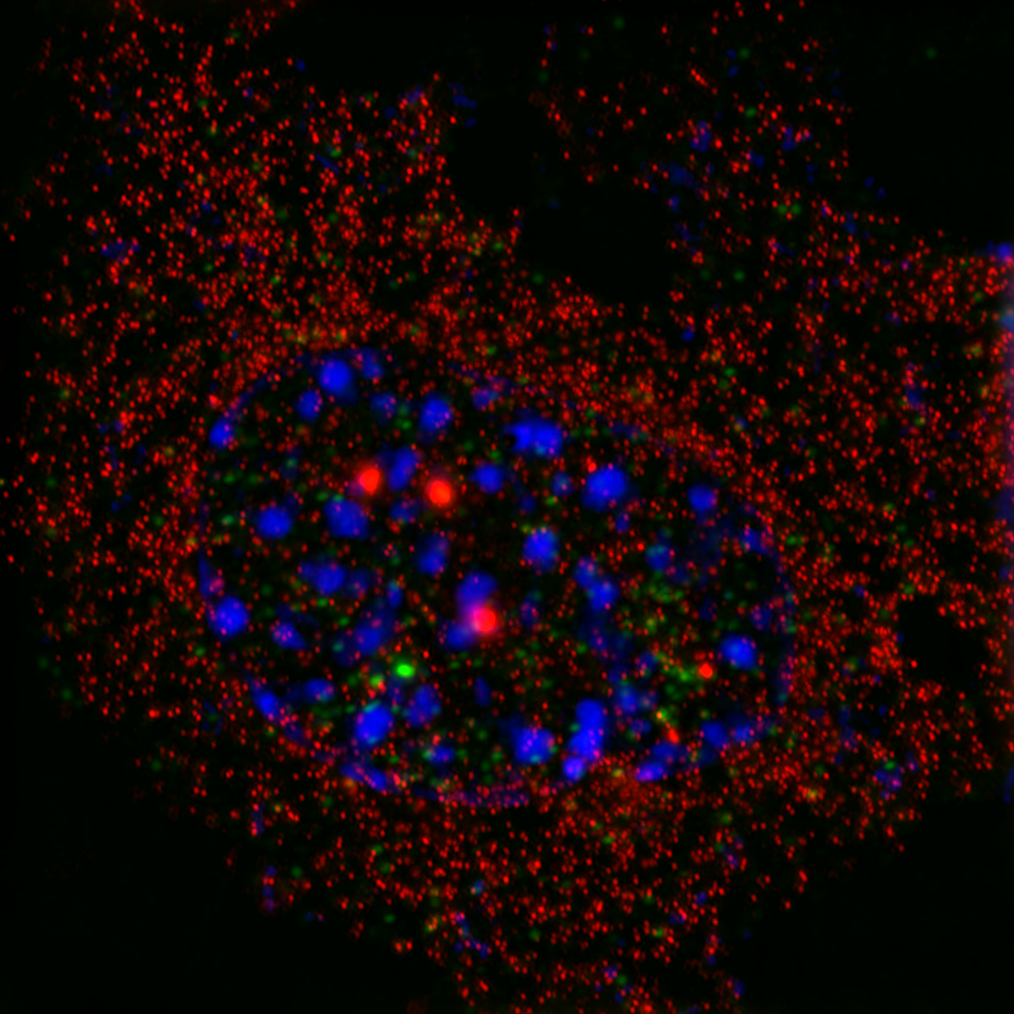
