## Supplementary material for "Injection molded open microfluidic well plate inserts for user-friendly coculture and microscopy": Monorail1 use protocol

### Monorail1: Device Protocols

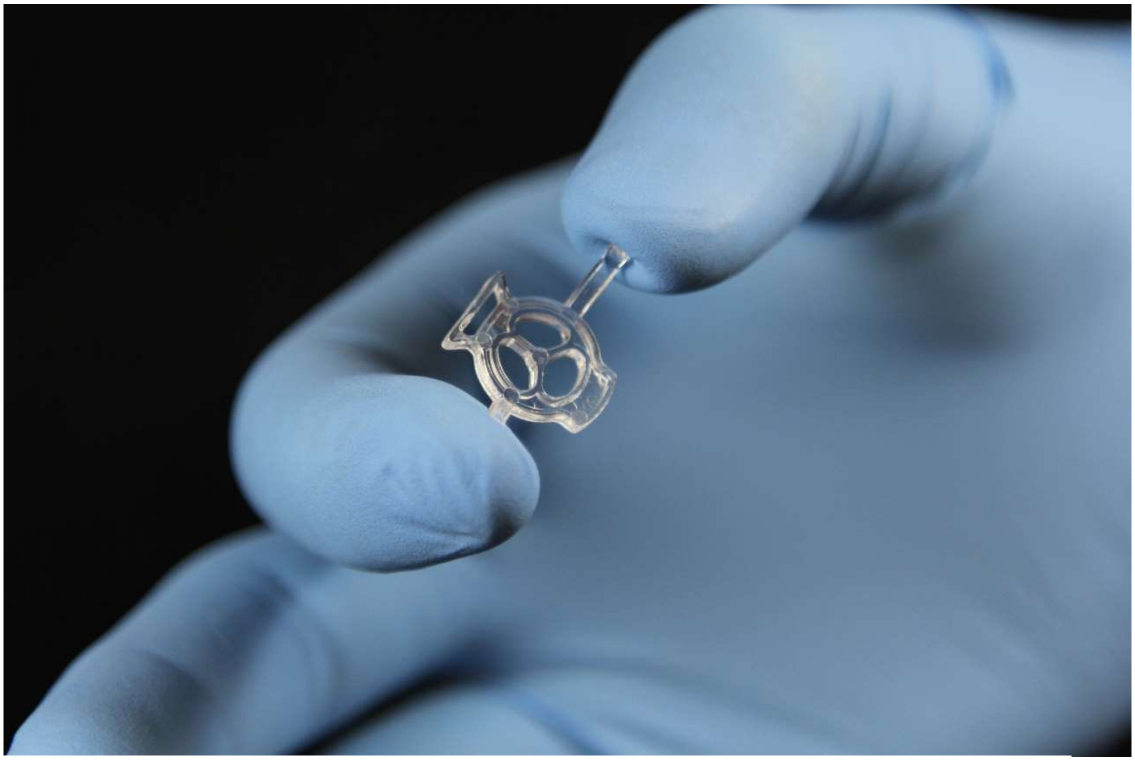

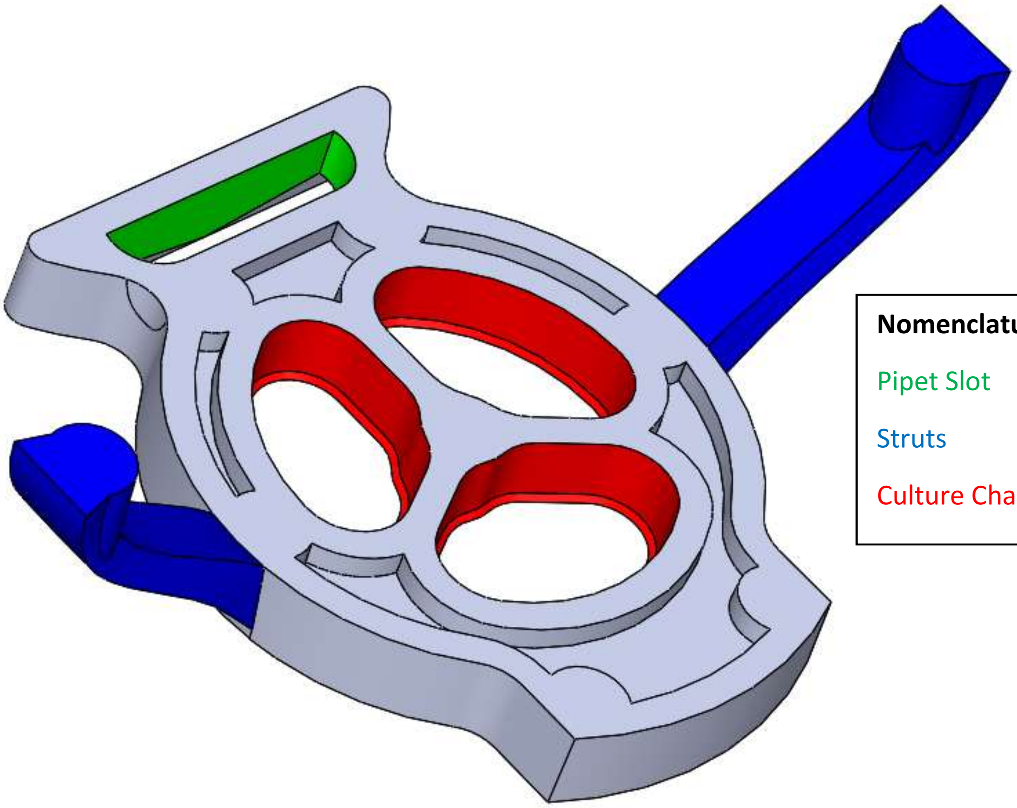

**Nomenclature Key:**

Pipet Slot

Struts

Culture Chambers

**Protocol A: Test your hands with Monorail1 devices.**

Before using the monorail device for cell culture, it is highly recommended that the user follow this protocol to practice using the device.

#### General notes:

- The monorail device uses gel to create patterned permeable walls that separate regions on the base of a well in a 12-well plate. Typically, this device is used to study soluble factor signaling, where different cell types are seeded into the separate culture regions and remain physically separate while exchanging secreted soluble factors through the gel wall. The plastic monorail device remains in the well during the full culture period.
- Refer to the image on the cover page for the key regions of this device including the pipet slot, culture chambers, and struts.
- Collagen type I and Matrigel are the two gels that have been validated for use with this platform. These gels need to be kept on ice to prevent premature gelation. This protocol describes the use of collagen I gel.
- Collagen I stock is acidic and needs to be neutralized for gelation. After adding HEPES buffer to the acidic collagen I stock (as described in the protocol below), it is recommended that the user pipet the solution up and down several times to homogenize the solution. Avoid making air bubbles in the collagen solution at this step, as air bubbles will persist in the solution and make it difficult to draw accurate volumes from the solution.
- Once the stock collagen solution is neutralized, it should be used within one hour, as the neutralized collagen will gel spontaneously, even on ice.

#### Materials provided:

- Monorail devices (Designation: Monorail1)
- Purple, green, and blue food coloring solutions for “testing your hands”
- 10X HEPES buffer
    - Note this 10X HEPES buffer has already been prepared for you and consists of 500 mM HEPES with 10x PBS, pH 7.6
- 1.5% low gelling temperature agarose solution in DI water or PBS (Sigma, Cat# A9414), sterilized by autoclave for cell culture use.

#### Additional materials required:

- 12-well plate
- Collagen solution (we recommend catalog number 354249 from Corning Inc. which is sold at a concentration ranging from 8-10.5 mg/mL depending on the lot)
- 1.5 mL or 0.5 mL centrifuge tube
- Needle nose tweezers

#### Device Protocol:

##### Fit devices into wells:

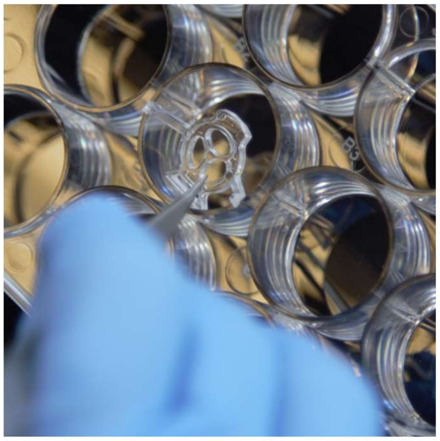

*Figure 1: Fit device into well.*

With a pair of needle-nose tweezers, grasp the monorail device and press it into the well of a 12-well plate, as seen in **Figure 1**. To ensure that the device is flush with the base of the well, use tweezers to press down on the device around the edges. **Figure 2** shows one such place where pressure should be applied, but it is advisable to also apply pressure in several other locations around the edge of the device.

##### Gel Loading:

##### Option 1: Collagen I

##### Prepare neutralized collagen gel solution:

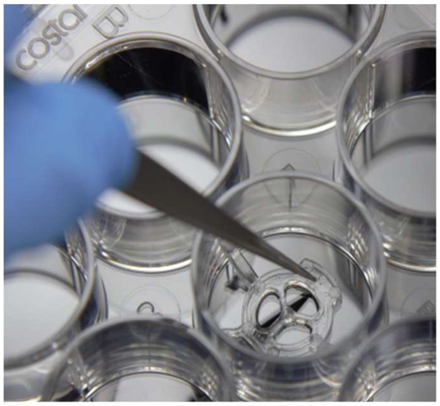

*Figure 2: Apply downward pressure to device.*

Keep the tubes containing collagen I on ice to prevent gelation. Prepare the neutralized gel solution by mixing 9 parts of collagen solution and 1 part of 10X HEPES in a microcentrifuge tube (mixing 36 µL stock collagen solution with 4 µL 10X HEPES is recommended per device).

##### Load neutralized collagen gel solution into device:

**Figure 3A** shows where neutralized gel solution (red) is pipetted into the device. *Note:* The collagen is dyed red in this image for visualization purposes for this protocol; do not add red dye to the collagen in normal experiments. Using a P200 micropipet, dispense 35 µL of neutralized gel solution into the pipet slot. **Figure 3B** shows the device photographed from under the well plate at four separate time points as collagen is following. Give the collagen about 10 seconds to flow under the entire device. Do not handle the well plate during this time, as tipping the well plate during flow can cause the device to fail. Once the flow is complete, incubate the well plate for 30 minutes. After this time interval, there should be a noticeable cloudiness in the neutralized collagen gel. This indicates that the collagen solution has gelled.

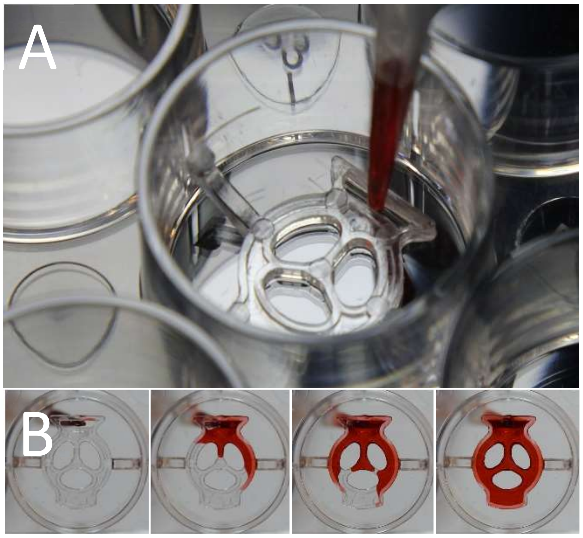


*Figure 3: Pipet neutralized collagen solution (red). A) Top view of pipet tip in pipet slot. B) Bottom view of device filling at 4 sequential time points.*

**Option 2: Low Gel Temp. Agarose**

Heat low gel temp. agarose solution using water bath or mini dry bath incubator to 50-55 °C. Well plates with devices prepared bay be maintained at room temperature in sterile conditions prior to hydrogel loading. Using a P200 micropipette, draw up 35 µL of low gel temp. agarose solution. As shown in **Figure 3A**, dispense 35 µL of low gel temp. agarose solution into the pipet slot with P200 micropipette. Give the low gel temp. agarose about 2 minutes to flow under the entire device. Do not handle the well plate during this time, as tipping the well plate during flow can cause the device to fail.

**Check for successful device loading:**

To ensure that the gel walls surrounding the culture chambers are intact, load 22 µL of food coloring into each of the 3 culture chambers. As in **Figure 4**, food coloring should be loaded directly into the center of each culture chamber. Using a different color for each chamber, you should observe *no immediate mixing* of color between the chambers (in a successful device transport of molecules through the gel wall occurs via diffusion, which takes time and will not be observed immediately). **Figure 5** shows the device photographed from above the well plate for a successful device (**Figure 5A**) and failed device (**Figure 5B**). *Note:* We recommend adding food coloring to the culture chambers for the purposes of this "test your hands" protocol only; food coloring is not normally added during actual cell culture experiments.

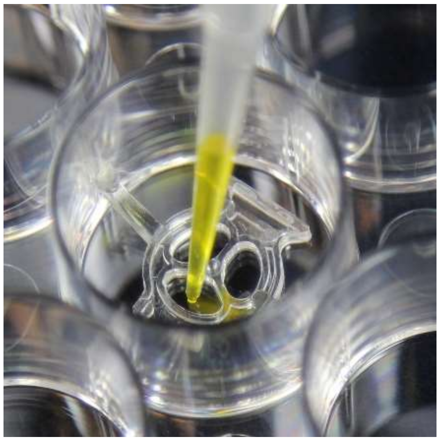

***Figure 4****: Load culture chambers with food coloring.*

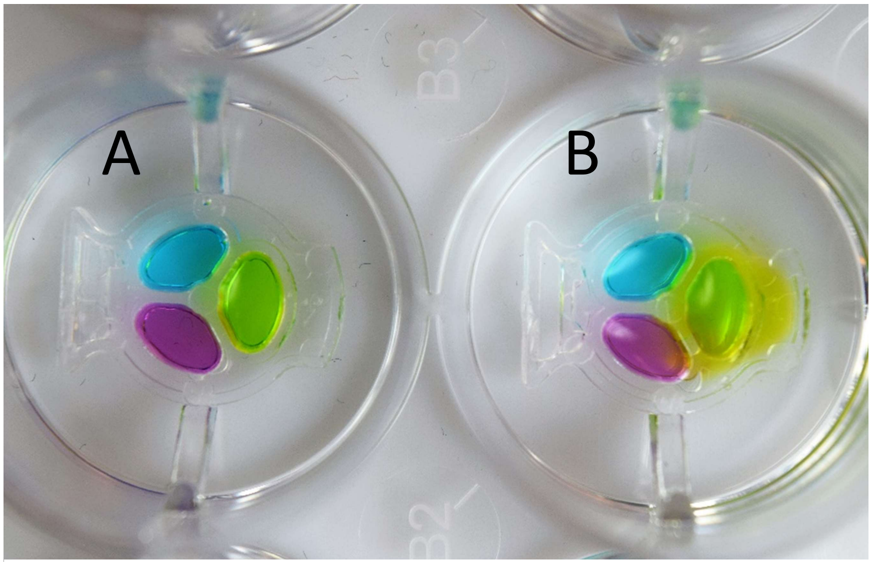


***Figure 5****: Two devices imaged after completion of protocol.* ***A)*** *A successful device where colors in each chamber (purple, green, and blue) do not mix.* ***B)*** *A failed device where green dye is seen rapidly dispersing outside of the culture chamber and some purple dye is seen in green chamber; both are due to incomplete filling of the gel wall.*

**Protocol B: Cell culture in Monorail1 devices.**

This document will outline the procedures required to pattern gel walls on either tissue culture treated surfaces or glass surfaces using monorail devices *(Designation: Monorail1)*. Before using this device, please read “Protocol A: Test your hands with monorail devices.”

#### Device specifications: Cell culture region surface areas and volumes

- Gel loading volume: 35 µL
- Culture area (each of the three culture chambers): 13.1 $mm^2$
- Culture volume (each of the three culture chambers): 22 µL
- Gel wall height: 250 µm

#### Device preparation:

- The following have already been done in the Theberge-Berthier lab:
    - Devices were sonicated for 60 min in isopropyl alcohol.
    - Devices were soaked for 30 min in 70% ethanol and air-dried.
    - The devices were plasma treated.
- The following must be completed by the user before using the monorail device for cell culture:
    - **If you are culturing cells on glass coverslips:**
        - Place sterile glass coverslips in the wells of a 12-well plate and do any necessary surface coating required for the cell type being cultured (e.g., poly-lysine coating for enhanced cell adhesion).
        - Aspirate coating solution and thoroughly wash glass coverslips in wells with sterile deionized water.
        - *Note:* It is important that any coating steps be done first. *Rationale:* Collagen I has been observed to retain and then leach coating materials such as poly-lysine, which at high concentrations can cause cell death. Therefore it is not recommended to add coating materials after the collagen gel wall is created.
    - UV sterilize monorail devices in biosafety cabinets, 10 min each side.
    - Put devices into 12-well plate as seen in **Figure 1 in Protocol A.** (*Note:* The maximum number of devices that can be used per 12-well plate is 8 devices. The four corner wells must be left empty for filling sacrificial water as described below.)
        - **If you are using glass coverslips**, make sure that the entire footprint of the device is enclosed in the coverslip.
    - Press down on the device with tweezers as seen in **Figure 2 in Protocol A**. Make sure the device fits tightly in the wells.

#### Gel Loading:

##### Option 2: Collagen I

- Keep the tubes containing collagen I on ice to prevent gelation. Prepare neutralized collagen solution by mixing 9 parts of collagen solution (~8-10.5 mg/mL collagen from Corning Inc.) and 1 part of 10X HEPES buffer* in a microcentrifuge tube (mixing 36 µL stock collagen solution with 4 µL 10X HEPES is recommended per device). [*10x HEPES buffer: 500 mM HEPES with 10x PBS, pH 7.6.]
- Load 35 µL of neutralized Col I to the devices through the loading port on each device (see **Figure 3 in Protocol A** for the location of the loading port).

##### Option 2: Low Gel Temp Agarose

- Heat low gel temp. agarose solution using water bath or mini dry bath incubator to 50-55 $^{o}$C.
- Well plates with devices prepared bay be maintained at room temperature in sterile conditions prior to hydrogel loading.
- Using a P200 micropipette, draw up 35 µL of low gel temp. agarose solution.
- Dispense 35 µL of low gel temp. agarose solution into the pipet slot with P200 micropipette.
- Do not handle the well plate during this time, as tipping the well plate during flow can cause the device to fail.

#### Filling sacrificial water:

*A warning:* Since the volume used in monorail devices is much lower than normal cell culture in well pates and flasks, evaporation is a serious problem that can cause media osmolarity changes and result in cell death. Sacrificial water is an important requirement and should be filled in the empty wells and the spaces in

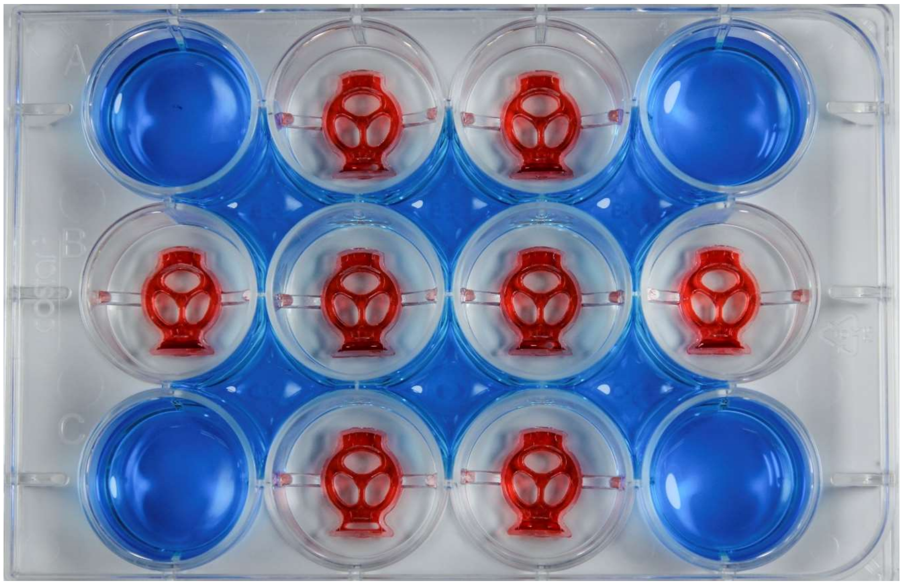

***Figure 6****: Sacrificial water (blue) placement in a 12-well plate. It is emphasized that sacrificial water must be added to all locations indicated. Color added for visualization purposes.*

between wells. As noted above, at a minimum the 4 corner wells of a 12-well plate need to be filled with sacrificial water, allowing 8 wells for devices as shown in **Figure 6 in this protocol (below)**. If fewer than 8 wells are used for devices, any additional empty wells should be filled with sacrificial water. Additionally, the well plate should be covered with the lid *immediately* after each manipulation.

- Add 2 mL of sterile water to each empty well and the spaces in between the wells in a 12-well plate as shown in **Figure 6**. (Here we have colored the sacrificial water blue for visualization.)
- Put the 12-well plate in 37 °C incubator and allow collagen to gel for 30-60 min.

**Seed cells into monorail devices:**

- Re-suspend cells in culture media at ~$3.1 \times 10^5$ cells/mL (this is the concentration we have used in test experiments with BHPrS-1 (stromal prostate), HLMVEC (endothelial cells) and MA10 (testis cells) cell lines; different concentrations can be used depending on the cell type of interest).
- Add 12 µL media to each culture chamber.
- Then add 10 µL cell suspension to each chamber to achieve a final seeding density of ~240 cells/$mm^2$ (again, this can be adjusted based on the desired seeding density for the cell type of interest).

**Maintaining cell culture:**

Due to the microscale volumes used with this device, cell culture media typically needs to be changed daily or more frequently (this may be more often than is required for conventional well pate culture). *Note:* Do not aspirate the media using vacuum as this can dislodge the device; rather gently aspirate the media with a pipette.
