## Supplementary material for "Injection molded open microfluidic well plate inserts for user-friendly coculture and microscopy": Monorail2 use protocol

### Monorail2: Device Protocols

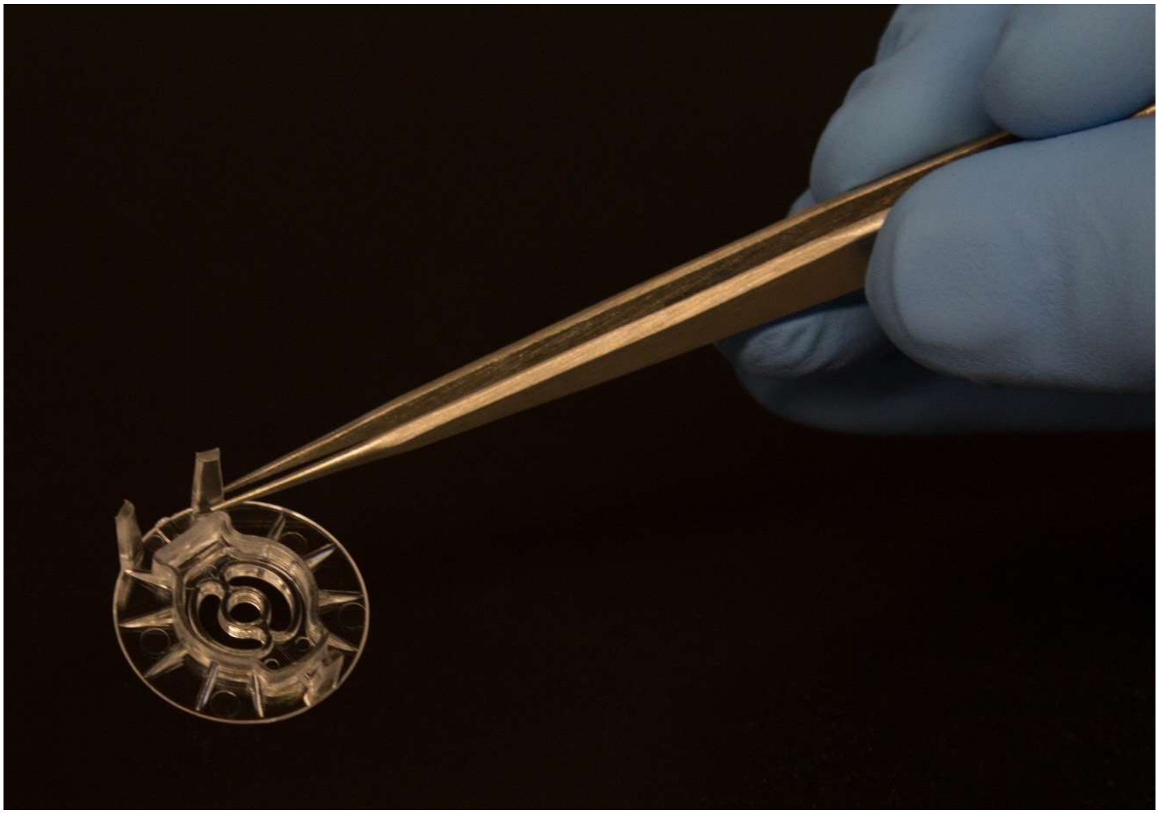

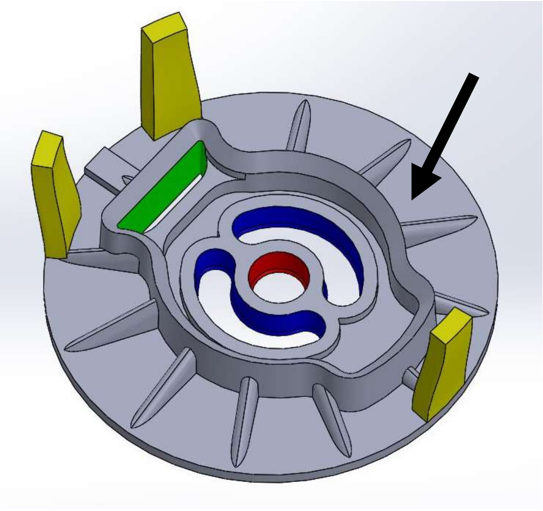

Nomenclature key:

Pressure Struts

Pipet Slot

Inner Cell Culture Chamber

Outer Cell Culture Chambers

**Arrow = Evaporation control outer ring**

**Protocol A: Test your hands with monorail devices.**

Before using the monorail device for cell culture, it is highly recommended that the user follows this protocol to practice using the device.

**General notes:**

- The monorail device uses gel to create patterned permeable walls that separate regions on the base of a well in a 12-well plate. Typically, this device is used to study soluble factor signaling, where different cell types are seeded into the separate culture regions and remain physically separate while exchanging secreted soluble factors through the gel wall. The plastic monorail device remains in the well during the full culture period.
- Refer to the image on the cover page for the key regions of this device including the pipet slot and culture chambers.
- Collagen type I, low gelling temperature agarose and Matrigel are gels that have been validated for use with this platform. Collagen I and Matrigel need to be kept on ice to prevent premature gelation. Low gelling temperature agarose needs to be heated to 50-55°C for use in this protocol. This protocol describes the use of collagen I gel andlow gelling temperature agarose.
- Collagen I stock is acidic and needs to be neutralized for gelation. After adding HEPES buffer to the acidic collagen I stock (as described in the protocol below), it is recommended that the user pipet the solution up and down several times to homogenize the solution. Avoid making air bubbles in the collagen solution at this step, as air bubbles will persist in the solution and make it difficult to draw accurate volumes from the solution.
- Once the stock collagen solution is neutralized, it should be used within one hour, as the neutralized collagen will gel spontaneously, even on ice.
- Agarose solution can be stored at room temperature. Prior to use for this protocol, it needs to be heated to 50-55°C.

**Materials provided:**

- Injection molded monorail devices
- 1:100 dilution food coloring solutions for “test your hands”
- 10X HEPES buffer
    - Note this 10X HEPES buffer has already been prepared for you and consists of 500 mM HEPES with 10x PBS, pH 7.6
- 1.5%low gelling temperature agarose solution in either DI H2) or PBS (Sigma, Cat# A9414), sterilized by autoclave for cell culture use.

**Additional materials required:**

- 12-well plate
- Collagen solution (we recommend catalog number 354249 from Corning Inc. which is sold at a concentration ranging from 8-10.5 mg/mL depending on the lot)
- 1.5 mL or 0.5 mL centrifuge tube
- Needle nose tweezers
- Water bath or mini dry bath incubator

### Preparing monorail devices:

#### Fit devices into wells:

1. With a pair of needle-nose tweezers, grasp the monorail device and press it into the well of a 12-well plate, as seen in **Figure 1**.
2. Press in multiple locations as shown with yellow arrows in **Figure 1**, to ensure the device is flat within the well.
3. **If you are using glass coverslips**, make sure that the entire footprint of the device is enclosed in the coverslip.

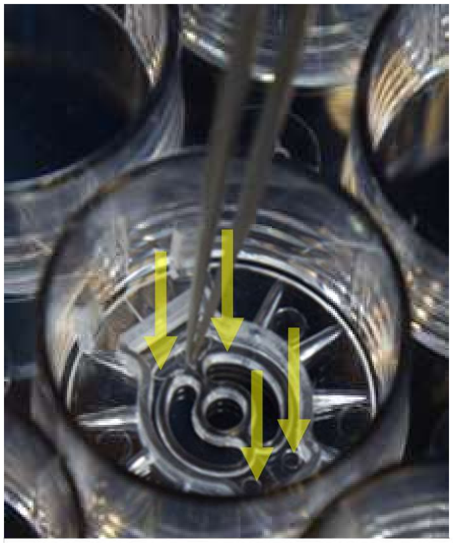

***Figure 1**: Load device into well plate. Ensure device is flat by pushing down in multiple places (indicated by yellow arrows).*

#### Gel Loading:

#### OPTION 1: Collagen I

Prepare neutralized collagen gel solution:

4. Keep the tubes containing collagen I on ice to prevent gelation.
5. Prepare the neutralized gel solution by mixing 9 parts of collagen solution and 1 part of 10X HEPES in a microcentrifuge tube. Due to collagen solution loss (in microcentrifuge tube and in pipet tip), prepare double the amount of neutralized gel solution than you expect to use.

Load neutralized collagen gel solution into device:

**Figure 2** shows where neutralized gel solution (red) is pipetted into the device. *Note:* The collagen is dyed red in this image for visualization purposes for this protocol; do not add red dye to the collagen in normal experiments.

6. Well plates with devices loaded may be maintained at room temperature in sterile conditions prior to hydrogel loading.
7. Using a P200 micropipette, very slowly draw up 40 µL of neutralized collagen solution.
8. As shown in **Figure 2**, Very slowly dispense 40 µL of neutralized collagen solution into the pipet slot with P200 micropipette.
9. Give the collagen about 2 minutes to flow under the entire device. Do not handle the well plate during this time, as tipping the well plate during flow can cause the device to fail.
10. If the collagen does not load or flow well into the device, do not use the device for your experiment.
11. Once the flow is complete, incubate the well plate for 30 minutes at 37°C.
12. After this time interval, there should be a noticeable cloudiness in the neutralized collagen gel. This indicates that the collagen solution has gelled.

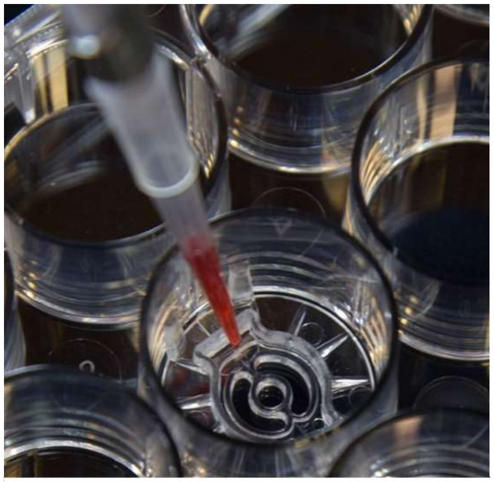

***Figure 2**: Load collagen into devices.*

**OPTION 2: Low gelling temperature agarose**

<u>Prepare low gelling temperature agarose:</u>

1. Prepare 1.5% low-gelling low gelling temperature agarose in DI $H_20$ or PBS. It will need to be sterilized with autoclave prior to use with cell culture. It can be prepared in advance and stored at room temperature.
2. Heat low gelling temperature agarose solution using water bath or mini dry bath incubator to 50-55°C.

<u>Load low gelling temperature agarose gel solution into device:</u>

3. Well plates with devices prepared may be maintained at room temperature in sterile conditions prior to hydrogel loading.
4. Using a P200 micropipette, draw up 40 µL of low gelling temperature agarose solution.
5. As shown in **Figure 2**, dispense 40 µL of low gelling temperature agarose solution into the pipet slot with P200 micropipette.
6. Give the low gelling temperature agarose about 2 minutes to flow under the entire device. Do not handle the well plate during this time, as tipping the well plate during flow can cause the device to fail.

**Check for successful device loading:**

**NOTE: Figures show triculture device, but test your hands is conceptually the same.**

7. Fill each inner culture chamber with 8 µL PBS or food coloring solution as shown in **Figure 3**. Take care to not dislodge the device with the pipet tip, as any sliding of the device could shear the gel wall from the bottom of the well, causing leakage.
8. Fill each outer culture chamber with 20 µL of a different color food coloring solution.
9. Load 500 µL of PBS or water into the area outside of the cell culture chambers (the evaporation control outer ring) by placing the pipet tip outside of the cell culture chambers, above the evaporation control ring (not pictured, see arrow on cover page).
10. You should observe *no immediate mixing* of color between the chambers or into the evaporation control outer ring (in a successful device transport of molecules through the gel wall occurs via diffusion, which takes time and will not be observed immediately).
11. Photograph devices immediately for analysis.
12. **Figure 4** shows the device photographed from above the well plate for a successful device (**Figure 4A**) and failed device (**Figure 4B**). *Note:* We recommend adding food coloring to the culture chambers for the purposes of this "test your hands" protocol only; food coloring is not normally added during actual cell culture experiments.

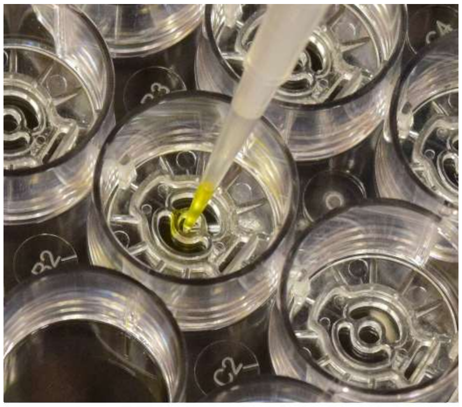

***Figure 3**: Load culture chambers with food coloring.*

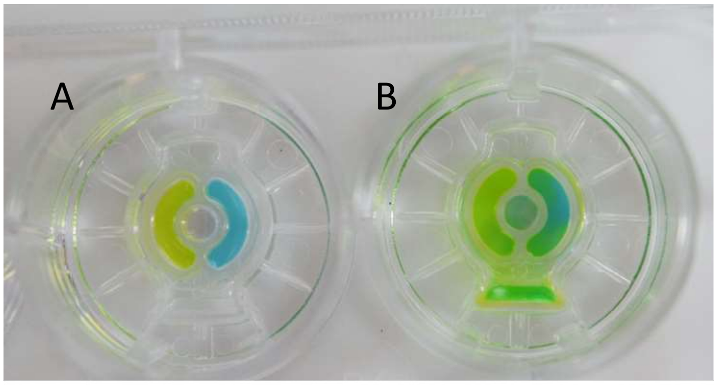


***Figure 4****: Two devices imaged after completion of protocol.* ***A)*** *A successful device where colors in each chamber (purple, green, and blue) do not mix.* ***B)*** *A failed device where green dye is seen rapidly dispersing outside of the culture chamber due to incomplete filling of the gel wall.*

**Protocol B: Cell culture in injection molded monorail devices.**

This document will outline the procedures required to pattern gel walls on either tissue culture treated surfaces or glass surfaces using monorail devices (specific design name: Monorail2).

**Device specifications: Cell culture region surface areas and volumes**

- Gel loading volume: 40 μL
- Center chamber culture area: 3.46 $mm^2$
- Outer chamber culture areas: 8.27 $mm^2$ (each outer chamber)
- Center culture volume: 8 μL
- Outer chamber culture volumes: 20 μL (each outer chamber)
- Gel wall height: 200 μm
- Sacrificial media volume: 500 μL

**Device preparation:**

- The following have already been done in the Theberge-Berthier lab:
  - Devices were sonicated for 60 min in isopropyl alcohol.
  - Devices were soaked for 30 min in 70% ethanol and air-dried.
  - The devices were plasma treated.

**Prior to using the monorail device for cell culture:**

1. UV sterilize monorail devices in biosafety cabinets, 10 min each side.
2. Plan your experiment:
   - We recommend a 4 or 8 device configuration (avoiding placement of devices in corners of 12-well plates), depending on your needs.
3. Coat well plates:
   - **If you are culturing cells on glass coverslips:**
     - Place sterile glass coverslips in the wells of a 12-well plate and do any necessary surface coating required for the cell type being cultured (e.g., poly-lysine coating for enhanced cell adhesion).
     - Aspirate coating solution and thoroughly wash glass coverslips in wells with sterile deionized water.
     - *Note:* It is important that any coating steps be done first. *Rationale:* Collagen I has been observed to retain and then leach coating materials such as poly-lysine, which at high concentrations can cause cell death. Therefore, we do not recommend adding coating materials after the collagen gel wall is created.
   - **If you choose to coat well plates with gelatin**, we recommend placing 2 mL 0.1% gelatin per well plate, incubate at 37°C for 30 minutes and then aspirate gelatin solution).

### Preparing monorail devices:

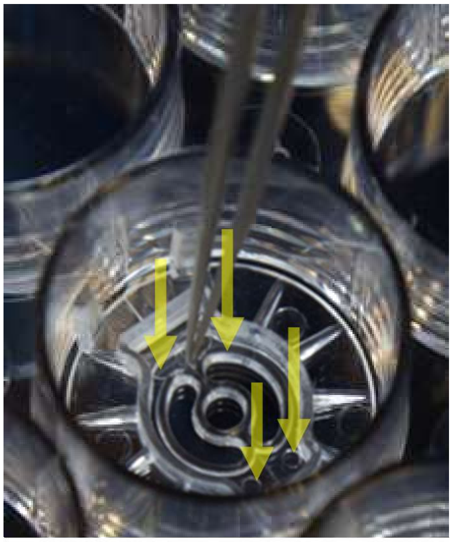

*Figure 1: Load device into well plate. Ensure device is flat by pushing down in multiple places (indicated by yellow arrows).*

**Fit devices into wells:**

1. With a pair of needle-nose tweezers, grasp the monorail device and press it into the well of a 12-well plate, as seen in **Figure 1**.
2. Press in multiple locations as shown with yellow arrows in **Figure 1**, to ensure the device is flat within the well.
3. **If you are using glass coverslips**, make sure that the entire footprint of the device is enclosed in the coverslip.

**Gel Loading:**

**OPTION 1: Collagen I**

Prepare neutralized collagen gel solution:

4. Keep the tubes containing collagen I on ice to prevent gelation.
5. Prepare the neutralized gel solution by mixing 9 parts of collagen solution and 1 part of 10X HEPES in a microcentrifuge tube. Due to collagen solution loss (in microcentrifuge tube and in pipet tip), prepare double the amount of neutralized gel solution than you expect to use.

Load neutralized collagen gel solution into device:

**Figure 2** shows where neutralized gel solution (red) is pipetted into the device. *Note:* The collagen is dyed red in this image for visualization purposes for this protocol; do not add red dye to the collagen in normal experiments.

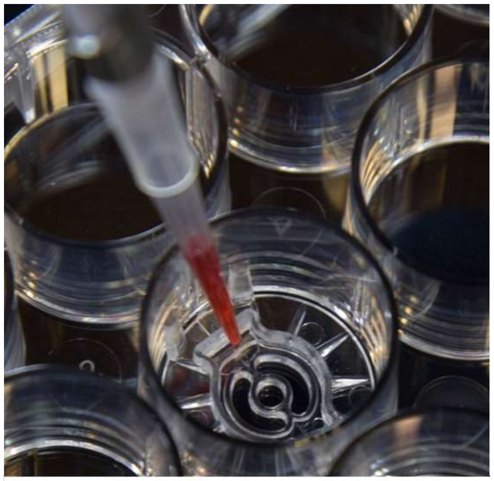

*Figure 2: Load collagen into devices.*

6. Well plates with devices prepared may be maintained at room temperature in sterile conditions prior to hydrogel loading.
7. Using a P200 micropipette, very slowly draw up 40 µL of neutralized collagen solution.
8. As shown in **Figure 2**, Very slowly dispense 40 µL of neutralized collagen solution into the pipet slot with P200 micropipette.
9. Give the collagen about 2 minutes to flow under the entire device. Do not handle the well plate during this time, as tipping the well plate during flow can cause the device to fail.
10. If the collagen does not load or flow well into the device, do not use the device for your experiment.
11. Once the flow is complete, incubate the well plate for 30 minutes at 37°C.
12. After this time interval, there should be a noticeable cloudiness in the neutralized collagen gel. This indicates that the collagen solution has gelled.

**OPTION 2: Low gelling temperature agarose**

Prepare low gelling temperature agarose:

13. Prepare 1.5% low gelling temperature agarose in DI $H_20$ or PBS. It will need to be sterilized with autoclave prior to use with cell culture. It can be prepared in advance and stored at room temperature.
14. Heat low gelling temperature agarose solution using water bath or mini dry bath incubator to 50-55°C.

Load low gelling temperature agarose gel solution into device:

15. Prepared well plates with devices may be maintained at room temperature in sterile conditions prior to hydrogel loading.
16. It is ok to hold the pipet containing the low gelling temperature agarose in your hand. Using a P200 micropipette, draw up 40 µL of low gelling temperature agarose solution.
17. As shown in **Figure 2**, dispense 40 µL of low gelling temperature agarose solution into the pipet slot with P200 micropipette.
18. Give the low gelling temperature agarose about 2 minutes to flow under the entire device. Do not handle the well plate during this time, as tipping the well plate during flow can cause the device to fail.

**Fill devices and/or seed cells**

At this point, you may proceed to seeding cells into device (described below), or in advance of planned cell seeding, fill each chamber with liquid (media or PBS).

19. Fill each inner culture chamber with 8 µL cell culture media or PBS
20. Fill each outer culture chamber with 20 µL cell culture media or PBS
21. Fill the chamber around each evaporation control ring with 500 µL cell culture media or PBS, (shown with black arrow on cover page).
22. Well plates may be stored at 4°C for up to two days before use for cell culture. Prior to use for cell culture, gently aspirate all liquid that is present and load fresh cell culture media as described in "Seed cells into monorail devices" section

**A note about evaporation control:**

These devices have been designed with an evaporation control ring, which forms an additional chamber inside the well for placement of sacrificial **media,** as seen in **Figure 3**. You must place 500 µL cell culture media in the evaporation control rings in order to successfully use the device. In other versions of our lab's monorail device, the user must add sacrificial water to the well plate corner wells and space between the wells, but this is not required in the current version of the device. It is still important that well plate should be covered with the lid *immediately* after each manipulation.

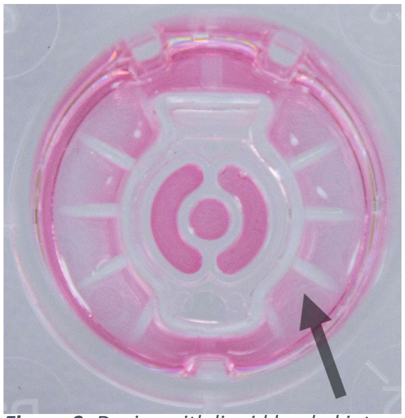

***Figure 3**: Device with liquid loaded into cell culture chambers and evaporation control ring (arrow).*

**Seed cells into monorail devices:**

23. If you have previously loaded devices with cell culture or PBS, gently aspirate all liquid that is present.
24. Add 4 µL media to the inner culture chamber.
25. Add 10 µL media to each outer culture chamber.

26. Add 500 µL media to the chamber formed by the evaporation control ring and the well (shown with arrow in Figure 3).
27. Re-suspend cells in culture media at ~$3.1x10^5$ cells/mL (this is the concentration we have used in a related device with BHPrS-1 (stromal prostate), HUVEC (endothelial cells) and MA10 (testis cells) cell lines; different concentrations can be used depending on the cell type of interest).
28. Add 4 µL cell suspension to the inner chamber to achieve a final seeding density of 250-300 cells/$mm^2$ (this can be adjusted based on the desired seeding density for the cell type of interest).
29. Add 10 µL cell suspension to each outer culture chamber to achieve a final seeding density of 250-300 cells/$mm^2$ (again, this can be adjusted based on the desired seeding density for the cell type of interest).

**<u>Maintaining cell culture:</u>**

Due to the microscale volumes used with this device, in cell culture chambers and evaporation control ring, cell culture media typically needs to be changed daily or more frequently (this may be more often than is required for conventional well plate culture). *Note:* Do not aspirate the media using vacuum as this can dislodge the device; rather, gently aspirate the media with a pipette.

**<u>Additional references of interest:</u>**

- An investigation of how to adjust culture conditions for microscale culture (in comparison to traditional macroscale well plate culture) to minimize cell stress, including considerations such as media changing frequency, increasing media buffering capacity, and adjusting cell seeding density:
  X. Su*, A. B. Theberge*, C. T. January, D. J. Beebe, "Effect of microculture on cell metabolism and biochemistry: Do cells get stressed in microchannels?" *Anal. Chem.* **2013**. *85*, 1562. (NCBI, ACS)
- Reference describing the original development of the monorail device:
  S. B. Berry*, T. Zhang*, J. H. Day, X. Su, I. Z. Wilson, E. Berthier, A. B. Theberge, "Upgrading well plates using open microfluidic patterning." *Lab Chip*, **2017.** *17*, 4253. (RSC)
